# An Inhibitory Function of TRPA1 Channels in TGF-β1-driven Fibroblast to Myofibroblast Differentiation

**DOI:** 10.1101/2022.04.12.488008

**Authors:** Fabienne Geiger, Sarah Zeitlmayr, Claudia A. Staab-Weijnitz, Suhasini Rajan, Andreas Breit, Thomas Gudermann, Alexander Dietrich

## Abstract

Transient receptor potential ankyrin 1 (TRPA1) is a non-selective Ca^2+^ permeable cation channel, which was originally cloned from human lung fibroblasts (HLFs). TRPA1-mediated Ca^2+^ entry is evoked by exposure to several chemicals, including allyl isothiocyanate (AITC), and a protective effect of TRPA1 activation in the development of cardiac fibrosis has been proposed. Yet, the function of TRPA1 in transforming growth factor β1 (TGF-β1)-driven fibroblast to myofibroblast differentiation and the development of pulmonary fibrosis remains elusive. TRPA1 expression and function was analyzed in cultured primary HLFs, and mRNA levels were significantly reduced after adding TGF-β1. Expression of genes encoding fibrosis markers, e.g. alpha smooth muscle actin (*ACTA2*), plasminogen activator inhibitor 1 (*SERPINE1*), fibronectin (*FN1*) and type I collagen (*COL1A1*) was increased after siRNA-mediated down-regulation of TRPA1-mRNA in HLFs. Moreover, AITC-induced Ca^2+^ entry in HLFs was decreased after TGF-β1 treatment and by application of TRPA1 siRNAs, while AITC treatment alone did not reduce cell viability or enhanced apoptosis. Cell barrier function increased after addition of TRPA1 siRNAs or TGF-β1 treatment. Most interestingly, AITC-induced TRPA1 activation augmented ERK1/2 phosphorylation, which might inhibit TGF-β-receptor signaling. Our results suggest an inhibitory function of TRPA1 channels in TGF-β1-driven fibroblast to myofibroblast differentiation. Therefore, activation of TRPA1 channels might be protective during the development of pulmonary fibrosis in patients.

## Introduction

Lung fibrosis is a chronic progressive disease with limited medical treatment options, leading to respiratory failure and death with a median overall survival time of 3-5 years after diagnosis (1). The most progressive form of the disease is idiopathic pulmonary fibrosis (IPF) (2), a chronic fibrotic interstitial lung disease of unknown cause, which occurs primarily in older adults (3). The incidence of IPF appears to be higher in North America and Europe (3 to 9 cases per 100,000 person-years) than in South America and East Asia (fewer than 4 cases per 100,000 person-years) (4).

The role of inflammation of lung tissues in IPF has been discussed extensively (5). However, multiple and potent anti-inflammatory therapies, in particular corticosteroids, have failed to show benefits in patients with IPF (5). The recently in the US approved drug pirfenidone down-regulates the production of growth factors like TGF-β1 and procollagens, whereas the already established nintedanib is a multi-tyrosine-kinase inhibitor. Both compounds slow down the decrease of forced vital capacity and also improve patients’ quality of life, but do not increase survival rates (6, 7). Therefore, the need for more causative acting drugs directed against novel identified pharmacological targets is obvious.

Although the detailed molecular steps of fibrosis development are still elusive, it is now generally thought that, after chronic microinjuries, epithelial cells release mediators like TGF-β1, which induce fibroblast to myofibroblast differentiation (1, 2). Myofibroblasts express more α-smooth muscle actin (α-SMA) and secrete extracellular matrix (ECM) proteins (e.g. collagen (COL), fibronectin 1 (FN1) and plasminogen-activator-inhibitor 1 (PAI-1)) for the recovery of lung barrier function (1, 2). Indeed, IPF is characterized by scattered accumulation of myofibroblasts in fibroproliferative foci with ECM, which results in irreversible destruction of lung architecture and seriously inhibits alveolar gas exchange in patients (1, 7). In addition to fibroblasts, alveolar epithelial cells (8, 9), fibrocytes (10), pericytes (11) and pleural mesothelial cells (12) may also, at least in part, transdifferentiate to myofibroblasts.

The superfamily of transient receptor potential (TRP) channels of vertebrates consists of 28 members in six families fulfilling multiple roles in the living organism (13). While TRPC6, the 6th member of the classical or canonical TRPC family, is activated by ligand binding (14), TRPV4, a channel of the vanilloid family, is thermosensitive in the range from 24 to 38 °C, and may serve as mechanosensor because it is activated by membrane and shear stretch (15). Both proteins form tetrameric unselective cation channels, which are expressed in human and mouse fibroblasts (16-18) and increase intracellular Ca^2+^ and Na^+^ levels initiating multiple signal transduction pathways and cellular responses (13). Most interestingly, TRPV4 has already been identified as an important player in pulmonary fibrosis in mice, and its expression was upregulated in lung fibroblasts derived from IPF patients (18). Moreover, we were able to show that mice lacking TRPC6 were partly protected from bleomycin-induced lung fibrosis (17). TRPC6 channel expression was increased in primary murine lung fibroblasts after application of TGF-β1, and ablation of TRPC6 resulted in reduced α-SMA production, less migration of myofibroblasts as well as in decreased secretion of ECM (17). Therefore, TRPV4 and TRPC6 are important determinants of TGF-β1-induced myofibroblast differentiation during fibrosis in mice.

Here, we set out to analyze TRP mRNA expression patterns in primary human lung fibroblasts (HLFs) with and without incubation with TGF-β1 in a more translational approach. While TRPC6 and TRPV4 mRNA expression was not changed, we identified a TGF-β1-mediated down-regulation of TRPA1, another member of the TRP superfamily. TRPA1, which is the only member of the TRPA family, harbors many ankyrin repeat domains in its amino-terminus and opens its pore after exposure to several chemicals including AITC (19). Most interestingly, steroids and pirfenidone, a drug used as therapeutical option in patients with lung fibrosis, also activated TRPA1 channels, which resulted in inhibition of trinitrobenzene-sulfonic-acid–induced colitis (20). Moreover, calcitonin-gene-related peptide (CGRP) production after activation of TRPA1 channels in cardiac fibroblasts worked as an endogenous suppressor of cardiac fibrosis (21).

SiRNA-mediated down-regulation of TRPA1 mRNA resulted in up-regulation of mRNA and protein expression of pro-fibrotic marker proteins. AITC-induced increases in intracellular Ca^2+^ levels [(Ca^2+^)_i_] were reduced and cell barrier function was increased after application of TRPA1 siRNAs. Moreover, TGF-β1-signaling was decreased by AITC-induced TRPA1 activation, probably by ERK phosphorylation and SMAD2 linker phosphorylation. Therefore, in contrast to TRPC6 and TRPV4, TRPA1 channels are able to inhibit fibroblast to myofibroblast differentiation and may serve as pharmacological targets in the development of new therapeutic options for pulmonary fibrosis.

## Material and Methods

### Cells

HLFs of healthy donors were obtained from Lonza (Basel, Switzerland, #CC-2512), PromoCell (Heidelberg, Germany, #C-12360) or from the CPC-M BioArchive for lung diseases at the Comprehensive Pneumology Center (CPC Munich, Germany). HLFs were used until passage 8. Ethical statements were provided by the suppliers.

### Transcriptomic Analysis by RNA-Seq

HLFs were incubated with TGF-β1 or solvent as described in the Data Supplement and RNA-seq was performed by IMGM Laboratories (Martinsried, Germany) on the Illumina NovaSeq® 6000 next generation sequencing system as described (22). Data for normalized counts and differential gene expression were obtained with the bioconductor package DESeq2 (23) and gene enrichment analysis with ClusterProfiler (24) in R version 4.1.2. The dataset has been deposited in the ArrayExpress database at EMBL-EBI (www.ebi.ac.uk/arrayexpress) under accession number E-MTAB-11629.

### Plasmin Assay

To indirectly quantify the secretion of PAI-1 from HLFs, 10 µl of the supernatant of HLFs, 10 µL of 10 mM of D-Val-Leu-Lys 7-amido-4-methylcoumarin (D-VLK-AMC) (in 10 % DMSO/H2O) (Sigma-Aldrich, #V3138) as plasmin substrate (25) and 80 µL TRIS/HCl (20 mM, pH7.4) were incubated for 4 h at 37 °C, 5 % CO2 in a black 96-wellplate with clear bottom. Fluorescence intensity of AMC was measured at λ_ex_ 380 nm; λ_em_ 460 nm using a TECAN plate reader (Tecan, Infinite M200 Pro). Higher fluorescence values indicate increased levels of plasmin processed by t-PA, as D-VLK-AMC is hydrolyzed by plasmin to AMC (26).

### Electroporation and Luciferase Reporter Assay

HLFs were electroporated with the Neon® Transfection system to achieve plasmid transfection as described (27). 250,000 HLFs were resuspended in 100 µL buffer R (Neon™ Transfection System 100 μL Kit, Invitrogen, MPK10096) and 4 µL of the Luciferase Reporter Plasmid (1 µg/µL pGL3-CAGA(9)-luc) containing the SMAD-sensitive part of the human PAI-1 promotor controlling the luciferase gene (28) was added. The electroporation was performed with 1400 Volt, 2 pulses à 20 ms.. HLFs were seeded in four wells of a 24-well plate in antibiotic free medium containing 2 % (Lonza, PromoCell) or 10 % (DZL) serum. After 24 h, the cells were treated with TGF-β1 (2 ng/mL) or solvent, medium contained 1 % Penicillin/ Streptomycin and 0.1 % serum. On the second day, the cells were pretreated with A-967079 (500 nM) or solvent for 30 min and stimulated with AITC (10 µM), JT010 (75 nM) or DMSO for 120 min. The fibroblasts were lysed in 50 µl lysis buffer per well (25 mM Tris/HCl pH7.4, 4 mM EGTA, 8 mM MgCl2, 1 mM DTT, 1 % Triton X-100) for 10 min. at RT. Luminescence was measured upon injection of 20 µl luciferase reporter assay substrate (Promega, #E1501) to each well using an OMEGA plate reader.

**All other Methods are described in the Data Supplement, which is available online**.

## Results

### Transcriptional and Functional Down-Regulation of TRPA1 Channels in Human Lung Fibroblasts (HLFs) after Application of TGF-β1

To investigate the role of TRP channels in TGF-β1-driven fibroblast to myofibroblast differentiation, we cultured primary HLFs from three human donors, applied TGF-β1 (2 ng/ml for 48 h) or solvent and extracted RNA for a thorough transcriptomic analysis. While mRNAs coding typical marker proteins for lung fibrosis and signaling pathways for extracellular matrix and structure organization were significantly up-regulated (*see* Figure E1A, B), none of the channels of the TRPC, TRPV and TRPM families showed significant changes in their mRNA expression levels after application of TGF-β1 (*see* Figure E1C-E). For TRPA1 mRNA, however, there was a clear trend for a TGF-β1-induced down-regulation (*see* Figure E1F), which turned out to be significant in a comparison of log2 fold changes (Table 1). Moreover, this significant reduction of TRPA1 mRNA after application of TGF-β1 was reproducible by repetitive quantitative RT-PCR experiments (Figure 1A). To successfully manipulate TRPA1 expression in lung fibroblasts, we tested an siRNA pool directed against TRPA1 mRNA and identified an 84,4 % down-regulation of TRPA1 mRNA, while a scrambled control siRNA showed no significant changes (Figure 1A). We tested four TRPA1 antisera, including two monoclonal antibodies reported to be selective for TRPA1 channels (29). However, in our hands these batches of antibodies showed no selectivity for the TRPA1 protein in HEK293 cells stably expressing the channel (*see* Figure E2). Therefore, TRPA1 function was analyzed by Ca^2+^ imaging of fibroblasts. Application of the specific TRPA1 activator, AITC, resulted in a transient increase of the intracellular Ca^2+^ concentration ((Ca^2+^)_i_), which was reduced in cells transfected with TRPA1 siRNAs (Figure 1B, C). Most interestingly, fibroblasts treated with TGF-β1 showed significantly reduced AITC-induced increases in (Ca^2+^)_i_ in comparison to control cells incubated with solvent only (Figure 1D, E). We obtained similar results with another specific activator of TRPA1 channels JT010 (30)(*see* Figure E3A-B). Therefore, TRPA1 expression and function is reduced after application of TGF-β1 to primary human fibroblasts. This AITC-induced increase in (Ca^2+^)_i_ was blocked by a specific TRPA1 inhibitor, A-967079 (A96), in the absence and presence of TGF-β1 (*see* Figure E3C, D).

**Table 1:** Log2 fold changes and *P*-values for TRP channel mRNA expression obtained by transcriptomic analysis of human lung fibroblasts treated with TGF-β1 or solvent. *P*-values < 0.05 are indicated in **bold**. n.a, not available

| TRP Channel | Log2 fold change | *P*-value | Adjusted *P*-value |
| --- | --- | --- | --- |
| TRPA1 | -1.649 | **0.003** | **0.047** |
| TRPC1 | -0.437 | 0.262 | 0.675 |
| TRPC2 | n. a. | n. a. | n. a. |
| TRPC3 | 0.173 | 0.842 | 0.969 |
| TRPC4 | -0.788 | 0.523 | 0.860 |
| TRPC5 | 0.528 | 0.807 | n. a. |
| TRPC6 | -0.205 | 0.728 | 0.939 |
| TRPC7 | n. a. | n. a. | n. a. |
| TRPM1 | -1.531 | 0.705 | n. a. |
| TRPM2 | 1.239 | 0.761 | n. a. |
| TRPM3 | -0.868 | 0.499 | n. a. |
| TRPM4 | 0.037 | 0.932 | 0.992 |
| TRPM5 | n. a. | n. a. | n. a. |
| TRPM6 | -0.793 | 0.700 | n. a. |
| TRPM7 | 0.001 | 0.997 | 1.000 |
| TRPM8 | -0.525 | 0.755 | n. a. |
| TRPV1 | -0.048 | 0.931 | 0.991 |
| TRPV2 | -0.618 | 0.433 | 0.809 |
| TRPV3 | 1.693 | 0.214 | n. a. |
| TRPV4 | 0.259 | 0.821 | 0.964 |
| TRPV5 | n. a. | n. a. | n. a. |
| TRPV6 | 1.239 | 0.761 | n. a. |

**Figure 1:**
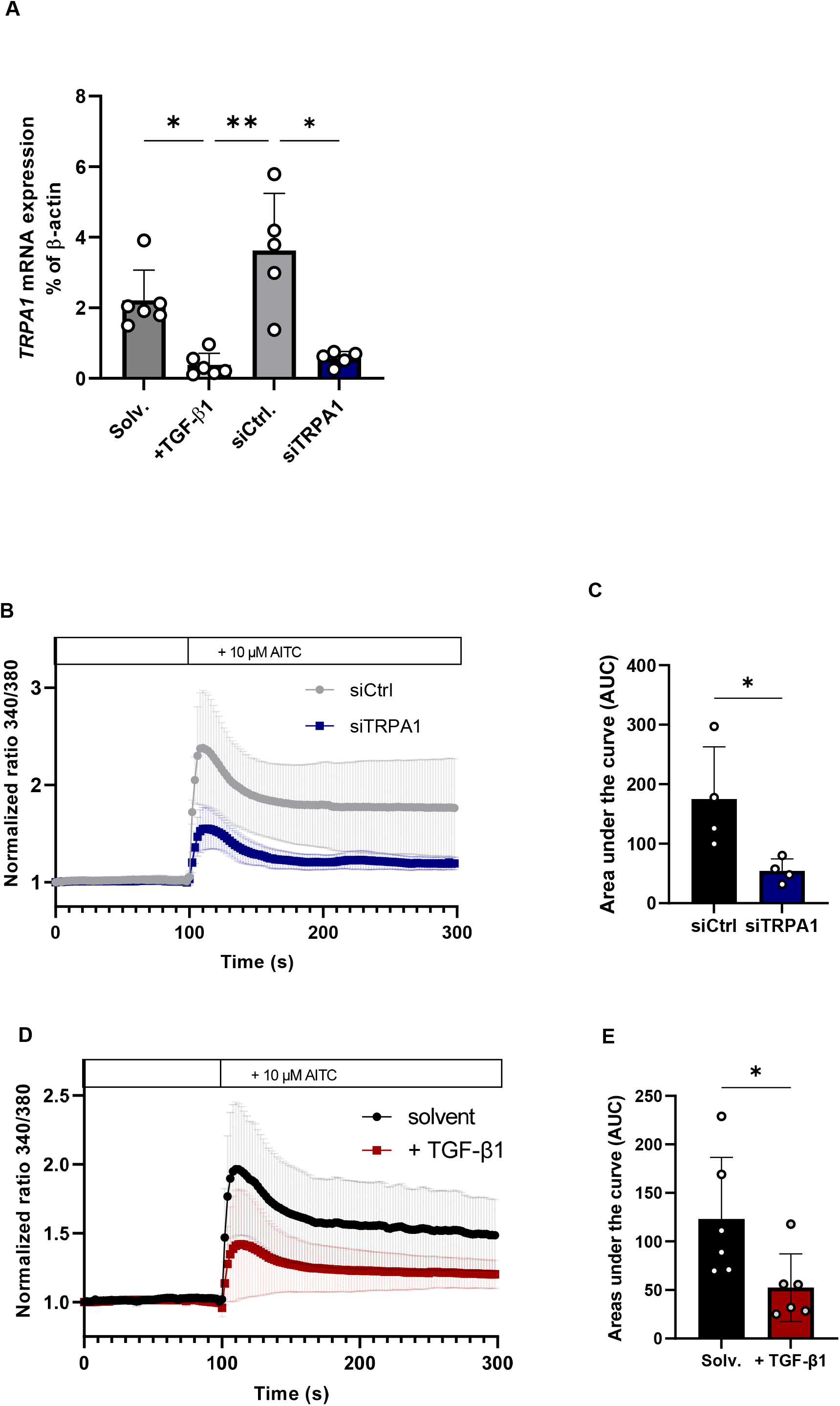
Quantitative reverse transcription (qRT)-PCR and Ca^2+^ imaging in primary human lung fibroblasts (HLFs) cultured with TGF-β1 or transfected with TRPA1-specfic siRNAs. (*A*) TRPA1 mRNA expression quantified by qRT-PCR experiments in HLFs cultured with TGF-β1 (+ TGF-β1) or transfected with TRPA1 specific siRNAs (siTRPA1). Solvent (Solv.) or scrambled siRNAs (siCtrl.) served as controls. (*B*) HLFs were transfected with TRPA1 specific siRNAs (siTRPA1, blue line) or scrambled siRNAs (siCtrl., grey line) as a control and intracellular Ca^2+^ levels were quantified as normalized ratios 340/380 nm after adding the TRPA1 activator AITC (10 µM) as described in Materials and Methods. Light grey areas represent SEM. (*C*) Areas under the curves were calculated and plotted as bars (siTRPA1, blue bar; siCtrl., black bar). (*D*) HLFs were cultured with TGF-β1 (+ TGF-β1, red line) or solvent (Solv., black line) as a control and intracellular Ca^2+^ levels were quantified as normalized ratios 340/380 nm after adding the TRPA1 activator AITC (10 µM) as described in the Data Supplement. Light grey areas represent SEM. (*E*) Areas under the curves were calculated and plotted as bars (+ TGF-β1, red bar; Solv., black bar). Data were analyzed by a Kruskal-Wallis (*A*) or a Mann-Whitney test (*C+E*) and are presented as mean ± SEM. qRT-PCR experiments were performed with n = 3 independent cell isolations each. For all graphs, *P < 0.05, ***P < 0,001, ****P <0.0001 versus the value in cells treated with solvent or scrambled siRNA alone.

### Down-Regulation of TRPA1 mRNA Increases Expression of Fibrosis Markers in HLFs

TGF-β1-induced fibroblast to myofibroblast differentiation is a key event in the development of lung fibrosis and results in up-regulation of gene expression of *ACTA2, COL1A1, FN1* and *SERPINE1*, encoding the fibrotic marker proteins α-smooth muscle actin (α-SMA), the α(1) chain of type I collagen (COL1A1), fibronectin 1 (FN-1) and plasminogen activator inhibitor 1 (PAI-1), respectively. As expected, application of TGF-β1 resulted in up-regulation of mRNA expression from these genes (Figure E1A). Most interestingly, transfection of cells by the specific TRPA1 siRNA but not a control siRNA was also able to increase mRNA expression of fibrotic marker proteins in the absence of TGF-β1 (Figure 2A-D). To evaluate these small but significant increases in mRNA levels on a protein level, we performed quantitative Western Blotting of protein lysates from fibroblasts transfected with TRPA1-specific or control siRNA. Expression of both proteins α-SMA and COL1A1increased after down-regulation of TRPA1 channels (Figure 2E-H). TRPA1-specific siRNA was however not effective, if cells were preincubated with TGF-β1 (see Figure E4A, B). Thus, in turn, TRPA1 expression and function seems to suppress expression of pro-fibrotic genes.

**Figure 2:**
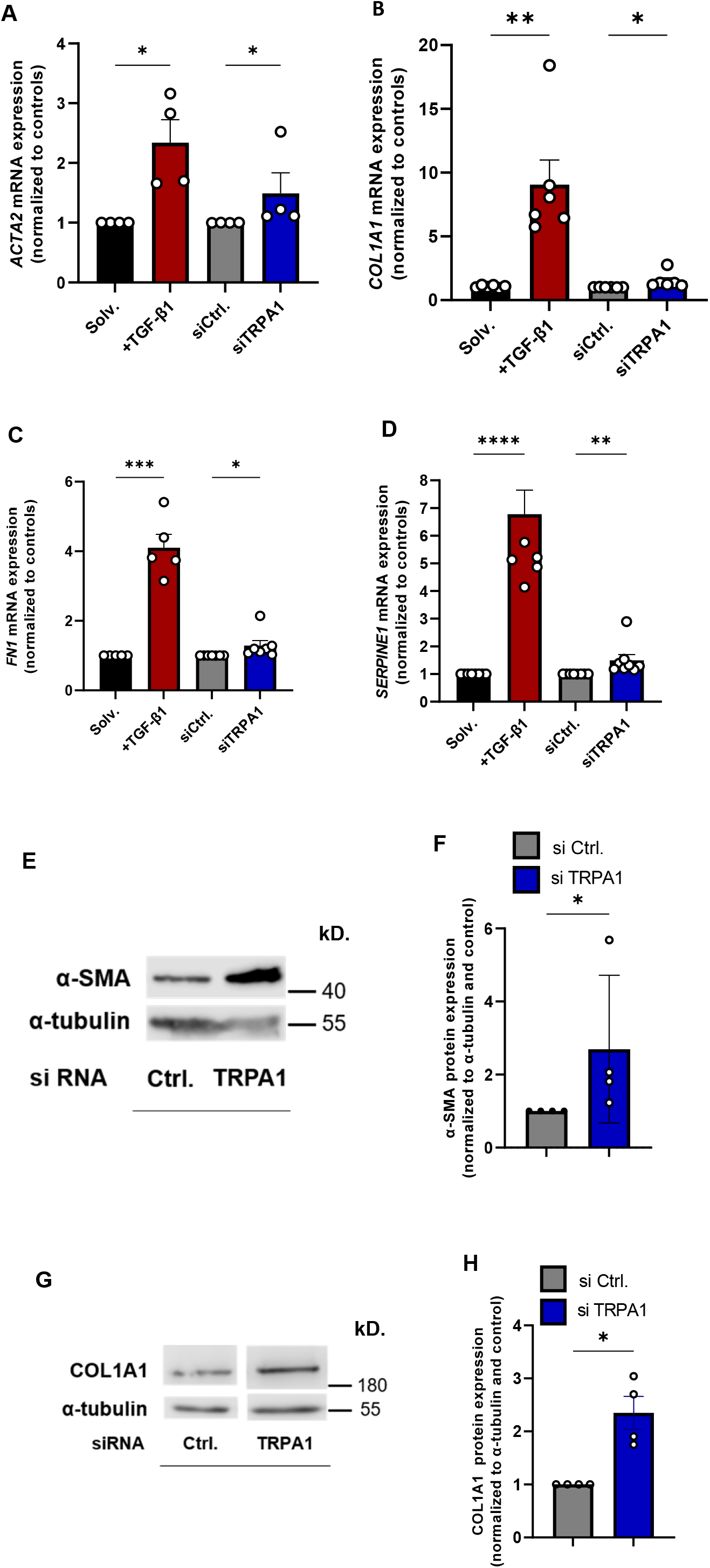
Quantitative reverse transcription (qRT)-PCR and Western-Blotting experiments for expression of fibrotic marker genes in human lung fibroblasts (HLFs) cultured with TGF-β1 or transfected with TRPA1-spefic siRNAs. (*A*) Quantification of mRNA levels of *α-SMA* (*ACTA2*) in HLFs cultured with TGF-β1 (+ TGF-β1) or solvent (Solv.) as well as transfected with TRPA1 specific siRNAs (siTRPA1) or scrambled siRNAs (siCtrl.) as control. (*B*) Quantification of mRNA levels of collagen 1A1 (*COL1A1*) in HLFs cultured with TGF-β1 (+ TGF-β1) or solvent (Solv.) as well as transfected with TRPA1 specific siRNAs (siTRPA1) or scrambled siRNAs (siCtrl.) as control. (*C*) Quantification of mRNA levels of fibronectin (*FN1*) in HLFs cultured with TGF-β1 (+ TGF-β1) or solvent (Solv.) as well as transfected with TRPA1 specific siRNAs (siTRPA1) or scrambled siRNAs (siCtrl.) as control. (*D*) Quantification of mRNA levels of plasminogen activator inhibitor (*SERPINE1*) in HLFs cultured with TGF-β1 (+ TGF-β1) or solvent (Solv.) as well as transfected with TRPA1 specific siRNAs (siTRPA1) or scrambled siRNAs (siCtrl.) as control. (*E, F*) Quantification of α-smooth muscle actin (α-SMA) protein levels in HLFs transfected with TRPA1 specific siRNAs (siRNA TRPA1) or scrambled siRNAs (siRNA Ctrl.) as control by quantitative Western-Blotting. (*G, H*) Quantification of collagen1A1 (COL1A1) protein levels in HLFs transfected with TRPA1 specific siRNAs (siRNA TRPA1) or scrambled siRNAs (siRNA Ctrl.) as control by quantitative Western-Blotting. Data were analyzed by a Kruskal-Wallis (*A-D*) or a Mann-Whitney test (*F, H*) and are presented as mean ± SEM. Cells are from at least three independent donors (n ≥ 4). For all graphs, *P < 0.05, **P < 0.01,***P < 0.001, ****P <0.0001 versus the value in cells treated with solvent or scrambled siRNA alone.

### Down-Regulation of TRPA1 mRNA Increase and Application of TRPA1 Activators Decrease Function of Pro-Fibrotic Proteins in HLFs

To further evaluate effects of TRPA1 channels on protein expression and function, we performed several assays. Detection of α-SMA by binding of an α-SMA-specific antibody followed by a fluorophore-coupled secondary antibody revealed increased protein levels after TGF-β1-induction in HLFs, which were also identified after application of the TRPA1 siRNA (Figure 3A, B). Phalloidin-staining to detect F-actin was also enhanced after incubation of cells with TGF-β1 or TRPA1 siRNA (Figure 3C, D). TGF-β1-induced up-regulation of α-SMA and collagen 1A1 (COL1A1) was detected by Western blotting in HLFs and was significantly down-regulated by the TRPA1 activator AITC (Figure 3E-H). We obtained similar results in the presence of the other specific TRPA1 activator JT010 and co-treatment with TGF-β1 (*see* Figure E4C-F). PAI-1 is able to inhibit tissue plasminogen activator (t-PA) or urokinase (uPA)-induced production of plasmin, which results in degradation of fibrin clots (Figure 4A). This process is quantified by cleaving of D-Val-Leu-Lys 7-amido-4-methylcoumarin (D-VLK-AMC) by plasmin and increased emission of fluorescence from free AMC (Figure 4B) (26). As expected, plasmin-induced fluorescence of the HLF medium was decreased after incubation of cells with TGF-β1. Transfection of cells with TRPA1 siRNA, was also able to decrease fluorescence, although to a lesser extent (Figure 4C). Application of the TRPA1 activators AITC (Figure 4D) or JT010 (*see* Figure E4G) also resulted in a lesser or no decrease in plasmin activity, respectively. In conclusion, down-regulation of TRPA1 results in higher expression and activity of pro-fibrotic marker proteins, while treatment of HLFs with TRPA1 activators resulted in decreased plasmin activity most probably by a reduced expression of PAI-1.

**Figure 3:**
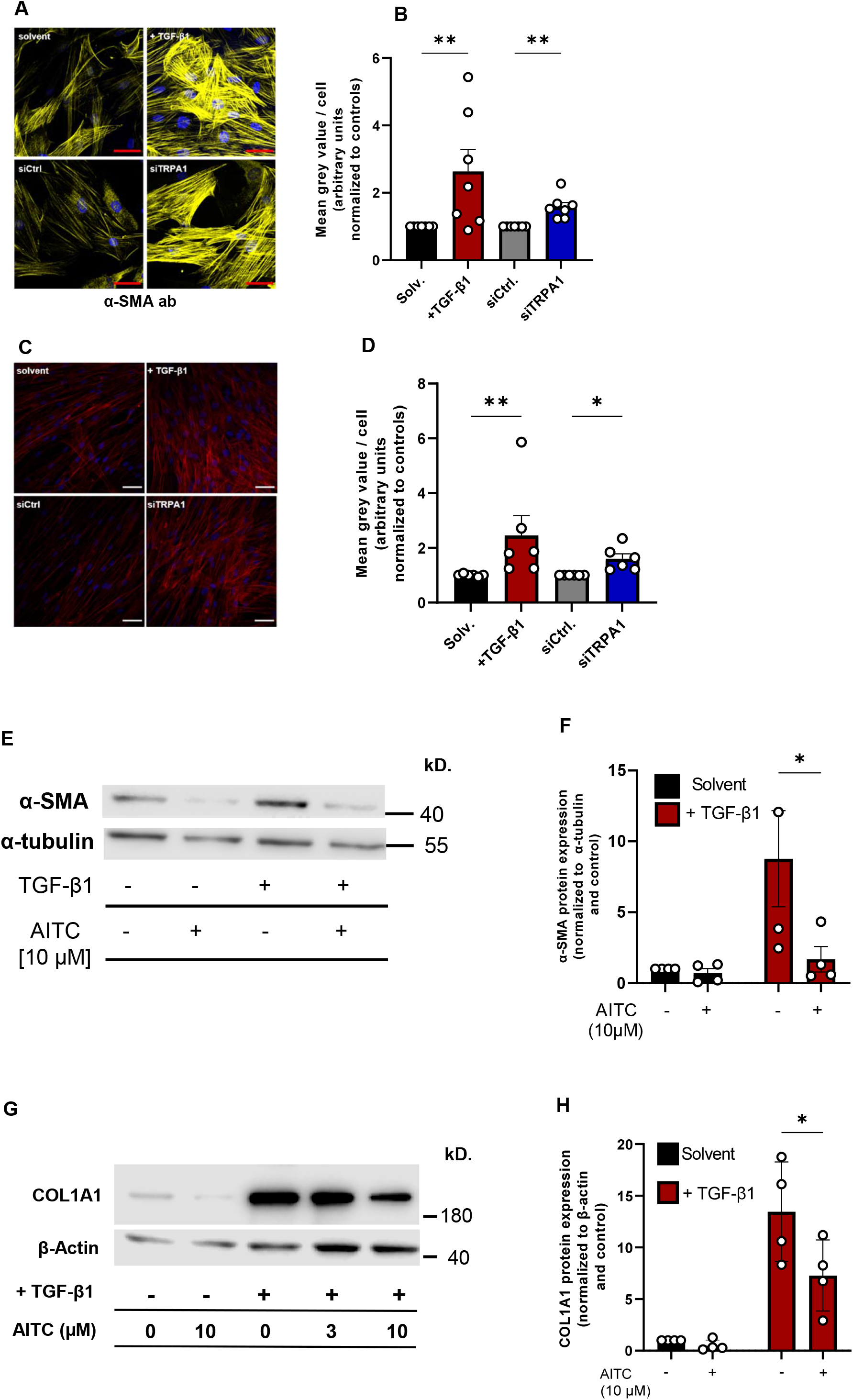
Quantification of expression of fibrotic marker proteins in human lung fibroblasts (HLFs) cultured with or without TGF-β1 and transfected with TRPA1-specific siRNAs or stimulated with the TRPA1 activator AITC. (*A*) Representative confocal laser microscopy images of HLF cultured with solvent (Solv.), TGF-β1 (+ TGF-β1), scrambled siRNA (siCtrl) or TRPA1-specific siRNA (siTRPA1) and stained with fluorophore-tagged anti α-smooth muscle actin antibodies (α-SMA ab). Scale bars, 50 µm. (*B*) Summary of mean gray values obtained from images shown in A. (*C*) Representative confocal laser microscopy images of HLFs cultured with solvent (Solv.), TGF-β1 (+ TGF-β1), scrambled siRNA (siCtrl) or TRPA1-specific siRNA (siTRPA1) and stained with fluorophore-tagged phalloidin for detection of F-actin. Scale bars, 50 µm. (*D*) Summary of mean gray values obtained from images shown in C. (*E*) α-smooth muscle actin (α-SMA) protein expression was quantified by Western blotting using a specific antiserum in HLFs cultured with or without TGF-β1 and/or treated with the TRPA1 activator AITC. Alpha-tubulin served as loading control. (*F*) Summary of α-SMA protein expression normalized to β-actin in HLFs cultured with or without TGF-β1 and/or treated with the TRPA1 activator AITC in the indicated concentrations. (*G*) Collagen 1A1 (COL1A1) protein expression was quantified by Western blotting using a specific antiserum in HLFs cultured with or without TGF-β1 and/or treated with the TRPA1 activator AITC in the indicated concentrations. Alpha-tubulin served as loading control. (*H*) Summary of collagen 1A1 (COL1A1) protein expression normalized to β-actin in HLFs cultured with or without TGF-β1 and/or treated with the TRPA1 activator AITC in the indicated concentrations. Data were analyzed by analyzed by a Kruskal-Wallis test (*B, D*) or two-way ANOVA (*F, H*) and are presented as mean ± SEM. Cells in A-F are from at least three independent donors (n = 4). For all graphs, *P < 0.05, **P < 0.01,***P < 0.001, ****P <0.0001 versus the value in cells treated with solvent or scrambled siRNA alone.

**Figure 4:**
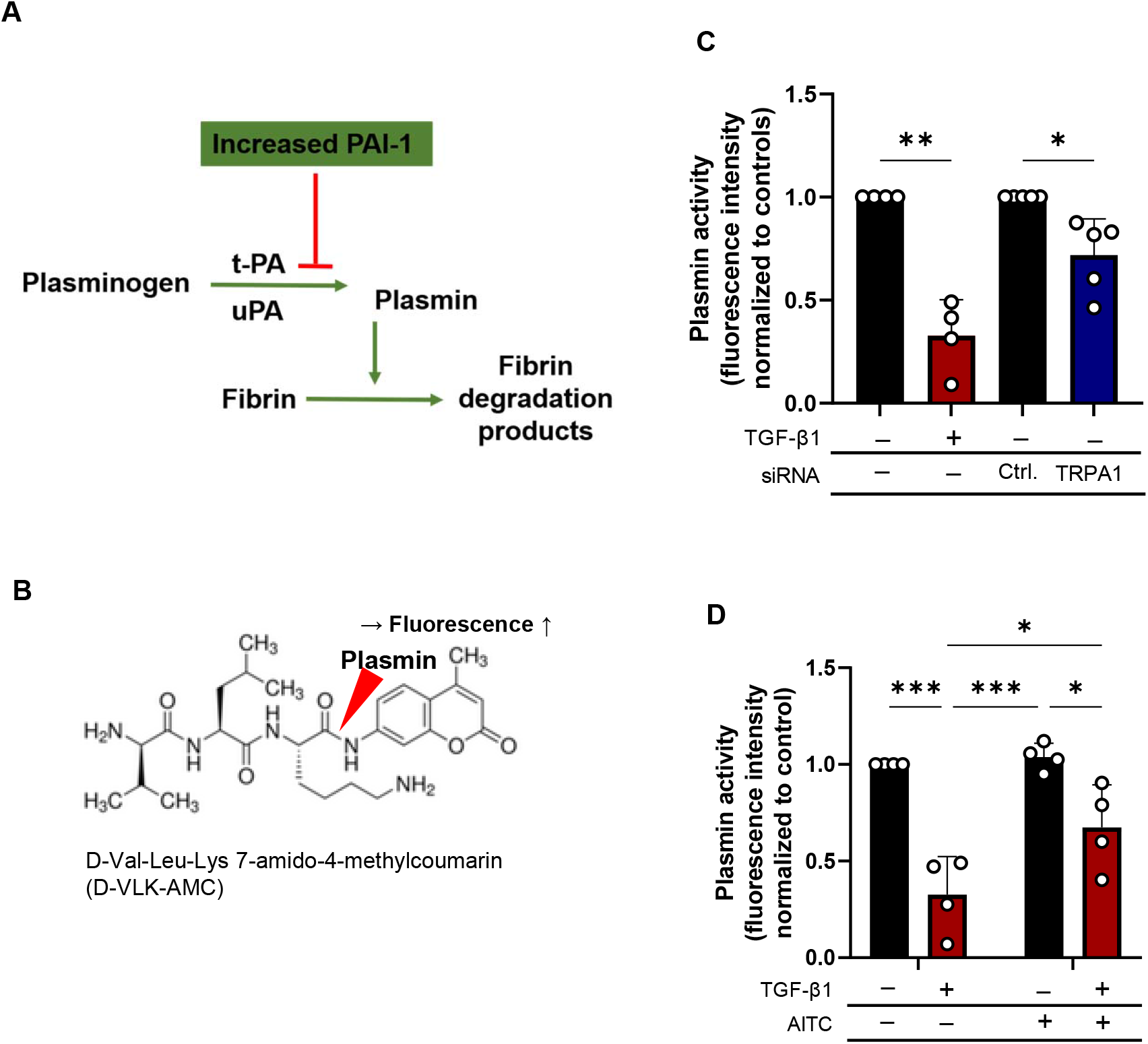
Indirect quantification of expression of plasminogen-activator-inhibitor 1 (PAI-1) activity in human lung fibroblasts (HLFs) cultured with TGF-β1 or transfected with TRPA1-specific siRNAs. (*A*) Schematic picture of PAI-1 inhibits cleavage of plasminogen in plasmin by tissue plasminogen activator (t-PA). Plasmin is able to degrade fibrin clots during fibrinolysis. (*B*) Hydrolysis of DVLK-AMC by t-PA results in free AMC, which is quantified by fluorometric detection (excitation 380 nm, emission 460 nm). (*C*) Plasmin activity quantified by AMC fluorescence in the medium of HLFs cultured with solvent (Solv.) or TGF-β1 (+ TGF-β1) and transfected with scrambled siRNA (siCtrl) or TRPA1-specific siRNA (siTRPA1). (*D*) Plasmin activity quantified by AMC fluorescence in the medium of HLFs cultured with (+) or without (-) TGF-β1 and with (+) or without application of the TRPA1 activator AITC. Data were analyzed by y a Kruskal-Wallis test (*C*) or two-way ANOVA (*D*) and are presented as mean ± SEM. Cells are from at least three independent donors (n = 3). For graph in C, **P < 0.01,***P < 0.001, ****P <0.0001 versus the value in cells treated with solvent or scrambled siRNA alone.

### Down-Regulation of TRPA1 mRNA Results in Increased Cell Resistance of HLFs

Lung fibrosis due to myofibroblast accumulation results in increased lung resistance. Along this line, we quantified cell barrier function in an ECIS system. Cell resistance increased after TGF-β1 treatment, most probably due to augmented α-SMA levels and production of ECM proteins like collagen and FN1 (Figure 5A, B). Most interestingly, application of TRPA1 siRNA to the extracellular medium also resulted in a significant increase in cell resistance (Figure 5C, D). In conclusion, down-regulation of TRPA1 mRNA changes cell barrier behavior of HLFs.

**Figure 5:**
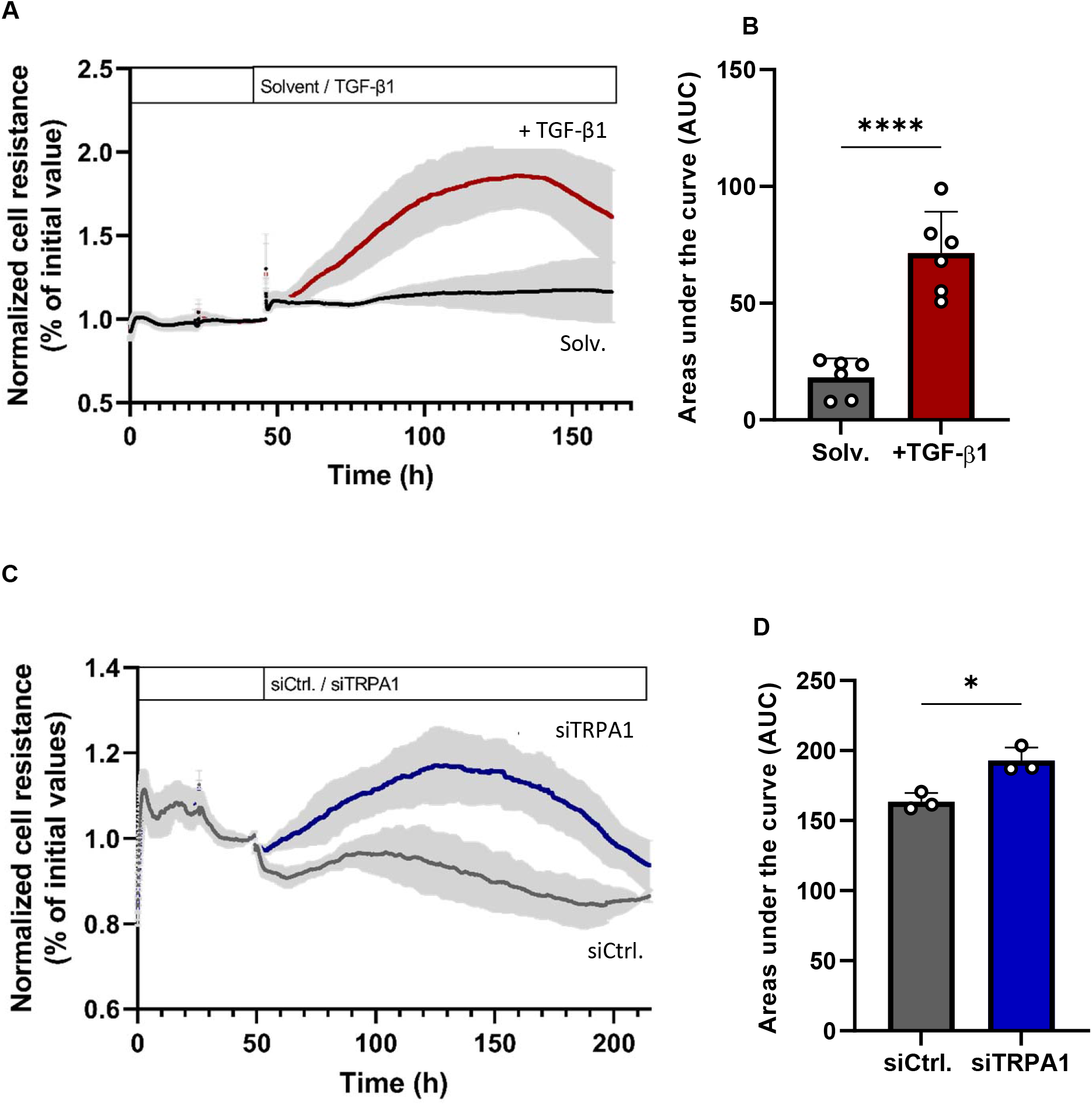
Electric cell-substrate impedance sensing in human lung fibroblasts (HLFs) cultured with TGF-β1 or transfected with TRPA1-spefic siRNAs. (*A*) Normalized cell resistance in % of initial values were quantified as described in the Data Supplement in HLFs cultured with TGF-β1 (+ TGF-β1) or solvent (Solv.). Light grey areas represent SEM. (*B*) Areas under the curves were calculated and plotted as bars (+ TGF-β1, red bar; Solv., grey bar). (*C*) Normalized cell resistance in % of initial values were quantified as described in the Data Supplement in HLFs transfected with TRPA1 specific siRNAs (siTRPA1, blue line) or scrambled siRNAs (siCtrl., grey line). Light grey areas represent SEM. (*D*) Areas under the curves were calculated and plotted as bars (siTRPA1, blue bar; siCtrl., grey bar). Data were analyzed by by an unpaired t test and are presented as mean ± SEM. Cells in A-D are from at least three independent donors (n = 3). For all graphs, *P < 0.05, ****P <0.0001 versus the value in cells treated with solvent or scrambled siRNA alone.

### TRPA1-Induced MAPK Phosphorylation Inhibits TGF-β1-mediated Transcription of Fibrotic Marker Proteins

To understand TRPA1 channel and TGF-β1 function on cellular transcription levels, we established a luciferase assay. A construct, containing luciferase cDNA under the control of *SERPINE1* (encoding PAI-1) core promoter region (pGL3-CAGA(9)-luc) (28), was transfected in HLFs. Cells were treated with TGF-β1, AITC, the TRPA1 inhibitor A-967079, or a combination of all three compounds (Figure 6A), and luciferase activity was quantified. As expected, application of TGF-β1 resulted in increased PAI-1 promoter-induced luciferase activity as compared to treatment with solvent only. Incubation of cells with AITC for 2 h, however, reduced TGF-β1-induced luciferase activity, which returned to initial levels if A-967079 was applied (Figure 6B). This inhibition of gene transcription was not due to cell apoptosis or reduced cell viability, as caspase and WST assays showed no increased apoptosis or reduced cell viability by AITC treatment, respectively (*see* Figure E7). TGF-β1 signaling via SMAD isoforms and inhibition of SMAD-dependent transcription by the MAPK signaling pathway has been reported in a fetal lung epithelial cell line (31, 32). Moreover, increased expression of pro-fibrotic markers by ERK inhibition was recently shown in dermal fibroblasts (33). Therefore, we quantified ERK1/2 and MAPK p38 phosphorylation in response to application of TGF-β1, AITC and TRPA1 siRNA by Western-analysis using a phospho-specific ERK1/2 antibody. AITC was able to increase ERK1/2 phosphorylation as reported before (34) as well as MAPK p38, which was reduced to basal levels after pre-incubation with TRPA1 siRNA (Figure 6C, D and Figure E5A-B). Pre-incubation with TGF-β1 resulted in ERK1/2 and MAPK p38 phosphorylation after application of TRPA1 siRNA or AITC, which however were not significantly different (see Figure E5C-F). TGF-β1 regulated transcription is mediated by carboxy-terminal phosphorylation of SMAD2/3 proteins, which are subsequently translocated to the nucleus. Additional phosphorylation of serine residues (245, 250, 255) in the linker region of SMAD2 proteins, however, results in inhibition of nuclear translocation ((31, 32) and reviewed in (35)). To dissect the signaling pathway mediating TRPA1-induced inhibition of TGF-β1 signaling in human fibroblasts, we analyzed linker phosphorylation of SMAD2 proteins in HLFs after stimulation by AITC by Ser245/250/255 specific phospho-SMAD2 antibodies. Most interestingly, application of AITC (3 µM) for 10 min. resulted in an increase in SMAD2 linker phosphorylation, which was found to be reduced after a longer incubation time of 30 min. and by simultaneous application of the TRPA1 inhibitor A567079 (Figure 6E, F). Preincubation with TGF-β1 also resulted in changes in SMAD2 linker phosphorylation after application of AITC, which however were not significantly different (see Figure E6A). Previous studies revealed that sustained stimulation of MRC5 cells or HLFs with TGF-β1 reduced total protein levels of SMAD3 (36, 37). We also observed a profound reduction of cytosolic SMAD3 levels after TGF-β1 stimulation of HLFs (*see* Fig. E6B, C). Interestingly, application of AITC significantly increased basal SMAD-3 levels (*see* Fig. E6B, C) confirming an important function of TRPA1 activation also on SMAD3.

**Figure 6:**
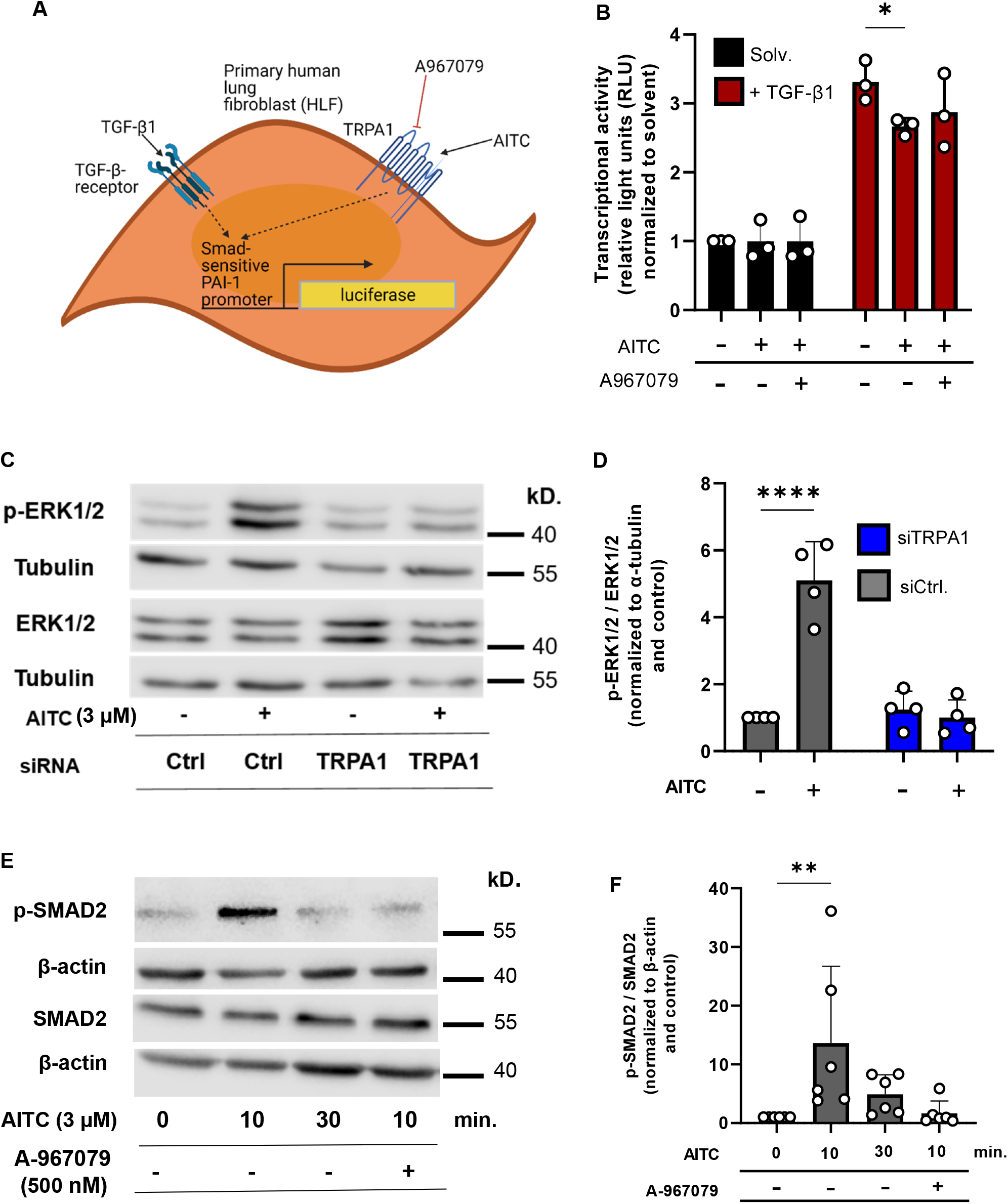
Identification of proteins involved in TRPA1-mediated inhibition of TGF-β1-induced transcription. (*A*) Schematic presentation of luciferase reporter assay. HLFs were transfected with a plasmid containing the luciferase gene under the control of the SMAD-sensitive part of the human PAI-1 promoter. TGF-β1 agonist for activation of SMAD transcription as well as AITC as TRPA1 activator or A-967079 as channel inhibitors were added and luciferase activity was quantified. (*B*) SMAD transcriptional activity in HLFs quantified after culturing in solvent (black bars) or TGF-β1 (red bars) as well as treatment with AITC or A-967079. (*C*) Lysates from fibroblasts transfected with TRPA1 specific (siRNA TRPA1) or scrambled siRNA (siRNA Ctrl.) and treated with (+) or without (-) AITC were incubated in a Western Blot of with specific antibodies directed against phosphorylated ERK1/2 (pERK1/2) and ERK1/2. Alpha-tubulin served as loading control. (*D*) Summary of the pERK1/2/ERK1/2 quantification normalized to alpha-tubulin by Western Blotting in HLFs transfected withTRPA1 specific siRNAs (siTRPA1, blue bars) or scrambled siRNAs (siCtrl., grey bars) incubated with (+) or without (-) AITC. (*E*) Lysates from fibroblasts pretreated with (+) or without (-) A-967079 and stimulated with (+) or without (-) AITC for the indicted times (min.) were incubated in a Western Blot of with specific antibodies directed against linker phosphorylated SMAD2 (pSMAD2) and SMAD2. Beta-actin served as loading control. (*F*) Summary of the pSMAD2/SMAD2 quantification normalized to β-actin by Western Blotting in HLFs pretreated with (+) or without (-) A-967079 and incubated with (+) or without (-) AITC. Data were analyzed by two-way ANOVA (*B, D*) or a Kruskal-Wallis test (*F*) and are presented as mean ± SEM. Cells in A-F are from at least three independent donors (n > 3). For all graphs, *P < 0.05, ****P <0.0001 versus the value in cells treated with solvent alone.

In conclusion, our data favor a mechanistic model in which TRPA1 mediated ERK phosphorylation inhibits TGF-β1-induced transcription and fibroblast to myofibroblast differentiation by linker phosphorylation of SMAD2 proteins (Figure 7).

**Figure 7:**
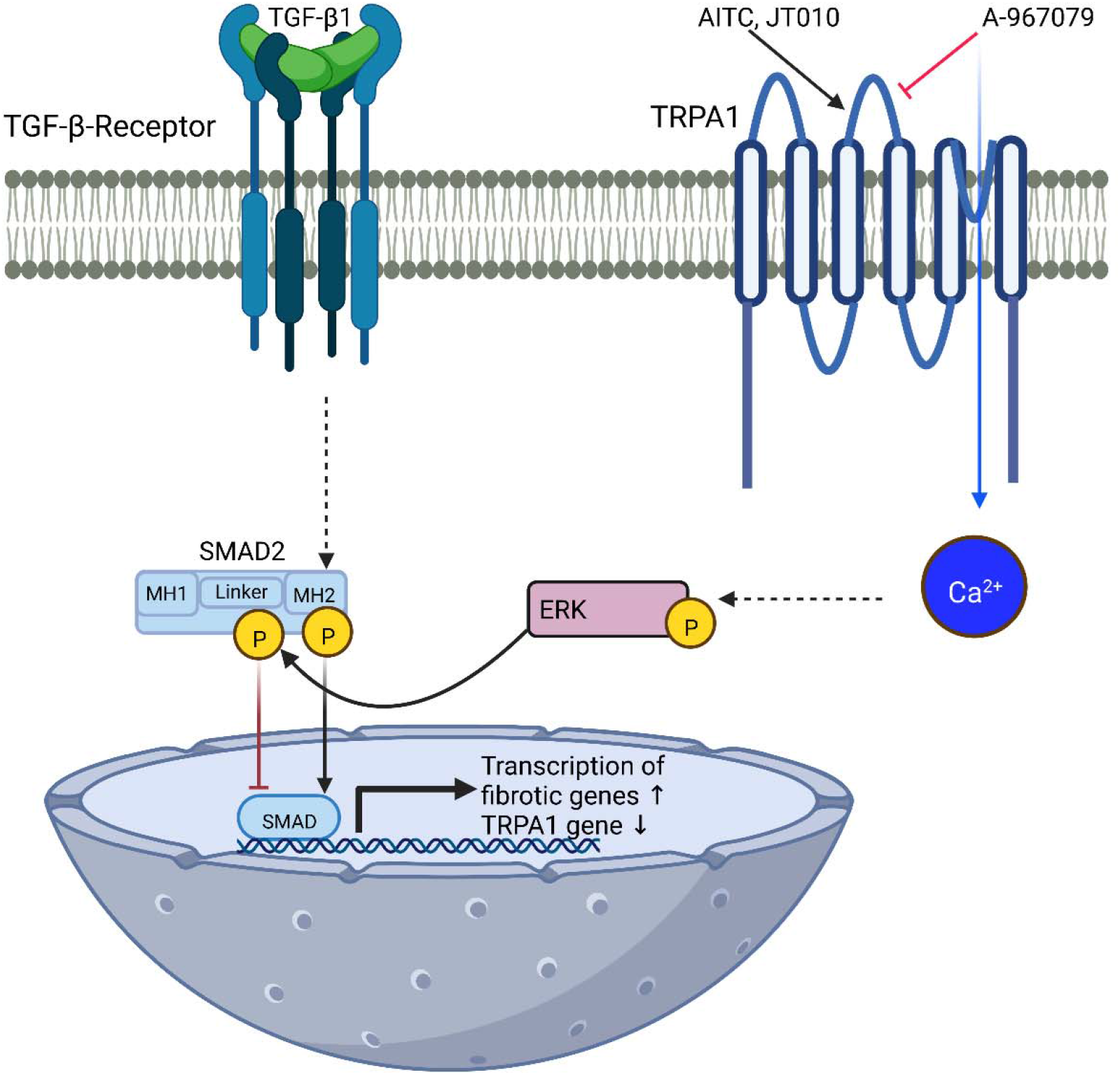
Proposed signaling pathway for TRPA1-mediated inhibition of TGF-β1-induced SMAD-mediated transcription. Ca^2+^ influx through AITC- or JT010 activated TRPA1 channels increases ERK1/2 phosphorylation, which induces linker phosphorylation of SMAD2. In contrast to the carboxyterminal phosphorylation of SMAD2, which enhances nuclear SMAD-induced transcription by activated TGF-β-receptors, linker phosphorylation of SMAD2 reduces TGF-β1-induced transcription of profibrotic genes. A TRPA1 inhibitor (A-967079) is able to reverse the inhibitory effect of TRPA1-induced Ca^2+^ influx. Moreover, TRPA1 gene expression is inhibited by TGF-β1-activation of HLFs by a still unknown mechanism.

## Discussion

Pulmonary fibrosis is a devastating disease with only a few treatment options. Although detailed molecular aspects of the development of lung fibrosis are still elusive, it is commonly believed that fibroblast to myofibroblast differentiation is a decisive step. The identification of new target proteins mediating this process is therefore an important step toward establishing new biomarkers and future therapeutic options. TRP-channels are expressed and essential for Ca^2+^ homeostasis in many cells including fibroblasts (17, 38). We were able to demonstrate an essential role of the classical TRP channel 6 (TRPC6) in murine fibroblast to myofibroblast differentiation (17). On a molecular level, TRPC6 mRNA and protein levels were up-regulated in myofibroblasts compared to fibroblasts and TRPC6 deficient mice were partially protected from bleomycin-induced pulmonary fibrosis (17). Moreover, expression of TRPV4, a member of the vanilloid family of TRP channels, was up-regulated in lung fibroblasts derived from IPF patients (18). However, after a thorough transcriptomic comparison of primary human fibroblasts with TGF-β1-differentiated myofibroblasts, we were not able to identify significantly different TRPC6 or TRPV4 mRNA expression levels. Most interestingly and contrary to mRNAs of all other TRP proteins, TRPA1 mRNA was found to be significantly down-regulated (Table 1). TRPA1 channels were originally identified in human fibroblasts (19). In addition, they are important sensor molecules for toxicants (39, 40). While the exact triggers for the development of lung fibrosis are still not known, it is believed that microinjuries of epithelial cells by chronic exposure to lung toxicants are an important predisposition for the disease (1, 7).

For the first time, we present evidence for an inhibitory role of TRPA1 channels in primary HLF to myofibroblast differentiation by TGF-β1. Preincubation of cells with TGF-β1 decreased AITC-induced increases in (Ca^2+^)_i_ (Figure 1D, E). A specific knockdown of TRPA1 expression increased mRNA (Figure 2A-D) and protein expression (Figure 2E-H) of important pro-fibrotic marker proteins, reduced AITC-induced increases in intracellular Ca^2+^ levels (Figure 1B-C) and increased cell resistance (Figure 5C-D). Most interestingly, we were able to monitor basal TRPA1 activity in HLFs, probably protecting these cells from myofibroblast differentiation. Of note, pre-application of TGF-β1 and TRPA1 siRNA had no additive effect on the expression of fibrotic genes (e.g. α-SMA and COL1A1 (Figure E4A, B), as TRPA1 mRNA might be already down-regulated in cells treated with TGF-β1. However, co-application of TGF-β1 and AITC or JT010 activated TRPA1 channels in the cell membrane with a high efficacy and reduced TGF-β1-mediated gene transcription and PAI-1 activity (Figure 6B, 4D and E4G) as well as expression of fibrotic proteins (e.g. COL1A1 and α-SMA in Figure 3E-H and E4C-F). On a molecular level, TRPA1 channel activation by AITC induced ERK1/2 phosphorylation (Figure 6C, D) and SMAD2 linker phosphorylation (Figure 6E, F), which reduces nuclear translocation and activation of transcription ((32) reviewed in (35)). Down-regulation by specific TRPA1 siRNAs or a TRPA1 inhibitor reversed these effects (Figure 6C-F). These findings highlight an inhibitory role of TRPA1 in the development of human myofibroblasts triggered by TGF-β1. In a recent publication, a similar effect was shown in MRC-5 and HF19 cell lines, which are both isolated from lungs of human fetuses and resemble some of the characteristics of human fibroblasts (41). TGF-β1 significantly down-regulated TRPA1 expression and AITC enhanced α-SMA gene and protein expression in these cell lines (41). HC-030031 as a TRPA1 antagonist however, failed to suppress the AITC-induced down-regulation of α-SMA and seemed to work synergistically with AITC (41). Only combined inhibition of ERK1/2 mitogen activated protein kinase (MAPK) and nuclear factor of erythroid 2-related factor (NERF) reversed the AITC-induced α-SMA suppression (41).

While this work was in progress, another research group investigated TRPA1 channels in human lung myofibroblasts (HLMFs) and TGF-β1-mediated pro-fibrotic responses (42). The researchers reported that the expression and function of TRPA1 channels in both healthy and IPF patients were reduced by application of TGF-β1 (42). TRPA1 over-expression or activation induced HLMF apoptosis, and TRPA1 activation by H_2_O_2_ induced necrosis. TRPA1 inhibition resulting from TGF-β1 down-regulation or pharmacological inhibition protected HLMFs from both apoptosis and necrosis (42). They concluded that TGF-β1 induces resistance of HLMFs to TRPA1 agonist- and H_2_O_2_-mediated cell death via down-regulation of TRPA1 (42). These data in human myofibroblasts confirm our results, underscoring an inhibitory role of TRPA1 in pro-fibrotic events governing fibroblast to myofibroblast differentiation at an early stage of lung fibrosis. However, there are also clear differences in the role of TRPA1 channels in HLFs versus HMLFs. We did not observe any reduction in cell viability or increases in apoptosis in human lung fibroblasts after activation of TRPA1 by the specific activator AITC (*see* Figure E7), while overexpression of TRPA1 in HLMFs increased apoptosis (42). However, we did not use H_2_O_2_, as this compound also activates TRPM2 channels and can induce cell death *per se* (43).

There is also evidence that TRP channels cluster with other signaling molecules in compartments formed by proteins like caveolins (44). In these caveolae as microdomains of calcium signaling (45), Ca^2+^ influx through TRPA1 channels might indeed inhibit fibroblast to myofibroblast differentiation, while in others Ca^2+^ influx through TRPV4 channels might promote lung fibrosis (18).

As it is generally accepted that fibroblasts mainly contribute to the progression of pulmonary fibrosis by differentiating to myofibroblasts and secreting excessive amounts of ECM (46), our data in HLFs and the results of others in HLMFs point to an important inhibitory function of TRPA1 channels in fibroblast to myofibroblast differentiation and HLMFs survival. Therefore, activation of TRPA1 channels may blaze the trail for new therapeutic options in lung fibrosis. Future studies will show, if TRPA1 activators are successful in fibrosis models, despite their effects on other cells (e.g. neurons), where TRPA1 channels are also expressed.

## Acknowledgement

This study was funded by the Deutsche Forschungsgemeinschaft (TRR152 (P15, P16) and GRK 2338 (P04, P07, P10)) and the German Lung Center (DZL). The authors thank Bettina Braun and Astrid Bauer for excellent technical assistance and Anna Litovskikh for her great help in the analysis of transcriptomic data with R. We gratefully acknowledge the provision of human primary fibroblasts from the CPC-M bioArchive and its partners at the Asklepios Biobank Gauting, the Klinikum der Universität München, and the Ludwig-Maximilians-Universität München. Figures 6A and 7 have been created by BioRender.com.

