## Supplementary material for "An Inhibitory Function of TRPA1 Channels in TGF-β1-driven Fibroblast to Myofibroblast Differentiation": Data Supplement

**Supplementary Materials and Methods:**

**Cell culture**

HLFs were grown in media containing 2 % (Lonza, #CC-3132; PromoCell, #C-23020) or 10 % (DZL (DMEM/F12, Lonza, #BE12-719F)) serum and seeded at a density of $3*10^5$ cells per well of a six well plate for experiments. On the next day, the medium was changed to starving conditions (0.1 % serum) or Accell siRNA delivery medium provided by the manufacturer (Dharmacon/HorizonDiscovery, Cambridge, UK, #B-005000-500) for 24 h, followed by rhTGF-β1 (bio-techne, Wiesbaden, Germany #240-B-002, 2 ng/ml) treatment for 48 h or siRNA knockdown (TRPA1 (#E-006109-00-0050)/scramble (#D-001910-10-50), 1 µM) for five days. A pool of four different TRPA1-specific siRNAs was used and siRNA transfection occurred without the use of transfection reagents by passive uptake of the siRNA. HLFs were pretreated with siRNA for three days and co-stimulated with siRNA and TGF-β1 for additional two days, if TGF-β1 treatment and TRPA1 knockdown were analyzed simultaneously. AITC (Sigma-Aldrich, Taufkirchen, Germany, #377430-5G, 3 µM or 10 µM as indicated) and the TRPA1 inhibitor A-967079 (Sigma-Aldrich, #SML0085-5MG, 500 nM) were applied as indicated.

**RNA-Seq**

Cell pellets were frozen in liquid nitrogen and stored at -80°C. RNA was isolated , followed by purification. The sequencing library was prepared with purified RNA with the Illumina TruSeq® Stranded mRNA technology. mRNA was enriched with a poly-T oligo pulldown. All the libraries were quality controlled by using DNA 1000/HS LabChip kits on the 2100 Bioanalyzer (Agilent Technologies). DNA was quantitfied with the highly sensitive fluorescent dye-based Qubit® ds DNA HS Assay Kit (Thermo Fisher Scientific). After pooling single libraries with an equal quantity of DNA, the sequencing library was finalized to 2.25nM and denatured with NaOH. Cluster generation and sequencing was performed on the Illumina® NovaSeq 6000. Real Time Analysis 3.4.4 Software (RTA) was used for primary image processing and bcl2fastq 2.20.0.422 software package for primary data analysis. Imaging and evaluation of the sequencing run performance was performed with the Illumina Sequence Analysis Viewer (SAV) 2.4.7. The reads were mapped to the human reference genome (GRCh38.p7) downloaded from NCBI.

**Quantitative Reverse-Transcription (qRT)-PCR**

Total RNA from HLFs was isolated using the RNeasy Plus Mini Kit (Qiagen, Hilden, Germany, #74136). The mRNA (1 µg) was transcribed into first-strand cDNA using RevertAid H MINUS 1$^{st}$ cDNA KIT containing reverse transcription polymerase (Life Technologies, Darmstadt, Germany, #K1631) and random primer according to the manufactures instructions. mRNA levels of target genes were analyzed by real-time quantitative PCR as previously described (1). 2 µl of the cDNA was added to 8 µl of a master mix consisting of 2x ABsolute QPCR SYBR Green Mix (Life Technologies,

#AB1158B), 10 pmol of each specific primer pair (see Table 1, Metabion, Planegg, Germany) and water. The following PCR program was run in a light-cycler apparatus (Roche, Mannheim, Germany): initial activation (15 min. at 94 °C) , 45 cycles of denaturation, annealing and extension (12 s at 94 °C, 30 s at 50 °C and 30 s at 72 °C) and melting curve determination to exclude samples that generated primer dimers or unspecific amplification products. The fluorescence intensities after each extension phase were measured, allowing the calculation of the crossing points (Cp) by the software (Roche, Basel, Switzerland, Light Cycler). Gene expression is shown in relation to a housekeeping control gene (*ACTB*) and the relative expression was determined using 2-ΔΔCt.

Table 1: DNA-Sequences of qRT-PCR primers

| Gene | Forward | Reverse |
|---|---|---|
| *ANKTM1* (*TRPA1*) (2) | TCACCATGAGCTAGCAGACTATTT | GAGAGCGTCCTTCAGAATCG |
| *ACTA2 (α-SMA)* | GAC CCT GAA GTA CCC GAT AGA AC | GGG CAA CAC GAA GCT CAT TG |
| *COL1A1* | CAA GAG GAA GGC CAA GTC GAG | TTG TCG CAG ACG CAG ATC C |
| *FN1* | CCG ACC AGA AGTTTGGGT TCT | CAATGCGGT ACA TGACCC CT |
| *SERPINE1 (PAI-1)* | GAC ATC CTG GAA CTG CCC TA | GGT CAT GTT GCC TTT CCA GT |
| *ACTB* | CCA ACC GCG AGA AGA TGA | CCA GAG GCG TAC AGG GAT AG |

**Western Blot**

Expression and phosphorylation of target proteins were evaluated by Western Blot analysis as previously described (1). HLFs were treated as described in section ”cells”, lysed with RIPA buffer (20 mM Tris-HCL, pH 7.5, 150 mM NaCl, 1 % Nonidet P40, 0.5 % sodium deoxycholate, 1 % SDS, 5 mM EDTA) containing phosphatase und protease inhibitors (Roche, Mannheim, Germany, #04906837001, #05892791001) for

30 min. on ice and sonicated for 15 s. The protein concentration was quantified using a Pierce$^{TM}$ BCA Protein Assay Kit (ThermoScientific, Rockford, US, #23225) following the manufacture's protocol. The protein samples (10 µg protein and 1x Laemmli (5x buffer: 3 ml TRIS/HCL (2.6 M), pH 6.8; 10 ml glycerin; 2 g SDS; 2 mg bromophenol blue and 5 ml β-mercaptoethanol)) were heated for 10 min. at 95 °C and loaded onto a 10 % SDS PAGE. Gel electrophoreses was performed for 30 min. at 70 Volts followed by 1.5 h at 100 Volts at RT. The proteins were transferred to a Roti®-PVDF membrane (Roth, Karlsruhe, Germany, #T830.1) using a wet blot system (BioRad, Feldkirchen, Germany) with a current of 250 mA for 1.5 h. The membrane was blocked with 5 % low fat milk (Roth, #T145.2) in TBS-T (0.1 %) for 1 h at RT. The antibodies were diluted in blocking solution and applied over night at 4 °C. HRP-conjugated secondary antibodies were applied for 2 h at RT. The chemiluminescence signal of the membranes was imaged after incubation in SuperSignal West Femto maximum sensitivity substrate (Life Technologies, #34095) with in an Odyssey-Fc-unit (Licor, Lincoln, NE, USA). Used antibodies and dilutions: p44/42 MAPK (ERK1/2) (Cell Signaling, #4695S, 1:1,000), Phospho-p44/42 MAPK (ERK1/2) (Thr202/Tyr204) (Cell Signaling, #4370S, 1:1,000), MAPK p38 (Cell Signaling #9212, 1:1000), pMAPK p38 (Cell Signaling #4511, 1:1000), α-smooth muscle actin (Sigma-Aldrich, #A5228, 1:2,000), Collagen1A1 (CellSignaling, #72026, 1:1,000), SMAD2 (CellSignaling, #3103 1:1,000), Phospho-SMAD2 (Ser245/250/255) (CellSignaling, #3104 1:1,000), β-actin-HRP (Sigma-Aldrich, #A3854, 1:10,000), α-tubulin (Abcam, #ab4074, 1:2,000), secondary anti-rabbit IgG peroxidase (POX)-antibody (Sigma-Aldrich, #A6154, 1:10,000), secondary anti-mouse IgG-HRP (CellSignaling, #7076S, 1:2,000).

**Electric cell-substrate impedance sensing (ECIS)**

The barrier function of cells can be quantified using electric cell-substrate impedance sensing (ECIS). After coating (L-Cysteine (10 mM, 10 min., RT) and FCS (overnight, 37 °C, 5 % $CO_2$)), HLFs were seeded at a density of 10,000 cells per well of a slide (ibidi, Gräfelfing, Germany, #8W10E+). A frequency of 8,000 Hz was applied and the cellular resistance was measured every two min. using an ECIS ZΘ (Applied Biophysics, Troy, NY, USA). Starving, TGF-β1/siRNA treatment and AITC stimulation were performed as mentioned above.

**Water Soluble Tetrazolium (WST) Assay**

To assess the cell viability of fibrotic and non-fibrotic fibroblasts after TRPA1 activation by AITC (3 µM, 24 h), HLFs were seeded at a density of 5,000 per well of a 24-well plate and treated. The cell culture media was aspirated, 500 µl of the cell proliferation WST-1 reagent (Sigma-Aldrich, #11644807001, 1:100 in medium,) was added to each well and incubated (37 °C, 5 % CO2, 3 h). The optical densities were measured in clear 96-well plates (100 µl/well in triplicates) using a plate reader (Tecan, Infinite M200 Pro).

**Caspase Assay**

Cell apoptosis was analyzed with the help of a CASPASE 3/7 Glo Assay (Promega, Walldorf, Germany, #G8091). HLFs were seeded at a density of 1,800 cells per well in a flat bottom white 96-well plate. siRNA (5 day), TGF-β1 (48 h) and AITC (24 h) treatment is described above. The assay was performed according to the manufactures protocol. In short, the appropriate amount of Caspase Glow reagent was added to the cells, media and reagent was thoroughly mixed (30 s, 300 rpm) and the assay was incubated for 1 h at RT. Luminescence was measured with a plate reader (Tecan, Infinite M200 Pro).

**Calcium Imaging**

TGF-β1/solvent or siRNA treated HLFs, which were grown on 25 mm coverslips until 80 % confluence, were loaded with Fura-2-AM (2 µM, Sigma-Aldrich, Taufkirchen, Germany, #47989-1MG-F) in $Ca^{2+}$-buffer (0.1 % BSA in HBSS (with $Ca^{2+}$, $Mg^{2+}$)/HEPES (0.5 M)) at RT for 30 min. HLFs were washed with $Ca^{2+}$-buffer and placed on the 40x oil-objective of a Leica DMi8 fluorescence microscope in a quick change chamber (Warner instruments, Holliston, USA, #64-0367) covered with 400/450 µl $Ca^{2+}$-buffer. The change of the intracellular calcium concentration after application of the specific TRPA1 agonists AITC (10 µM/3 µM) or JT010 (75 nM) with/out pre-blocking of the channel with A-967079 (500 nM) was measured at 340 and 380 nm as described (1).

**Immunofluorescence**

The cells were grown on 10 mm coverslips (three per six well) and treated as described in the section "cells" above. The HLFs were washed with PBS (Sigma-Aldrich, #D8537), fixed with 4 % PFA (Merck Millipore, Darmstadt, Germany, #104003) for 10 min at RT, permeabilized (0.5 % Triton X-100, 10 min, RT) and blocked with 4 % goat serum in 4 % BSA/PBS (1 h, RT). The primary antibody was applied over night at 4 °C in a wet chamber. The secondary antibodies were applied for 2 h at RT. Afterwards the cells were washed and the nuclei were stained using Hoechst (ThermoScientific, #62249, 2 µg/ml, 10 min, RT). The cover slips were fixed on glass slides using DAKO Fluorescence mounting medium (Agilent Technologies, Glostrup, Denmark, #S3023). Images were acquired at λex 488 nm; λem 525 nm with a confocal scanning microscope (LSM 880, Carl Zeiss). Antibodies and dilutions: αSMA (Sigma-Aldrich, #A5228, 1:2,000), Fibronectin (Abcam, #ab2413, 1:200), secondary anti-mouse IgG-FITC (Sigma-Aldrich, #F9006, 1:80), secondary anti-rabbit IgG-Alexa Fluor 488 (Invitrogen, #A11008, 1:500).

For the Phalloidin (Sigma-Aldrich, P1951) staining, the cells were treated as previously mentioned. Phalloidin was applied at a concentration of 50 µg/ml for 40 min at RT. The coverslips were mounted to the glass slides and images were taken at λex 540-545 nm; λem 570-573 nm using a confocal scanning microscope (LSM 880, Carl Zeiss). Image analysis was performed with the ImageJ-Fiji software calculating the mean grey values of the cells.


**Supplementary Figures:**

**Supplementary Figure Legends:**

**Figure E1:** Transcriptomic analysis of primary human lung fibroblasts (HLFs) after application of TGF-β1 or solvent. (*A*) Volcano plot of differentially expressed genes in TGF-β1 treated HLF Significantly up- (green color) or down-regulated red color) as well as fibrotic marker genes are marked: *ACTA2*, α-smooth muscle actin; *COL1A1*, collagen 1A1; *FN1*, fibronectin-1; *SERPINE1*, plasminogen activator inhibitor 1. (*B*) Significantly up-regulated gene ontology (GO) biological processes after application of TGF-β1. Adjusted *P*-values (p-adjust) are color-coded from blue to red in increasing order. Normalized counts of *TRPC* genes (*C*), *TRPM* genes (*D*) *TRPV* genes (*E*) and the *TRPA1* gene (*F*) in human fibroblasts after application of solvent (black bars) or TGF-β1 (red bars). Normalized counts were generated using DESeq2 in R (n = 3 independent cell isolations each). Data were analyzed by a Kruskal-Wallis (*C-E*) or a Mann-Whitney test (*F*) and are presented as mean ± SEM (n = 3).

**Figure E2:** Evaluation of the specificity of commercially available antibodies directed against the TRPA1 protein in HEK293 cell line stably expressing TRPA1 channels. (*A*) A representative $Ca^{2+}$ imaging experiment showing elevations of the intracellular $Ca^{2+}$ concentrations quantified as normalized ratios 340/380 nm after adding the TRPA1 activator AITC (10 μM) to HEK293 cells stably expressing TRPA1 protein (HEK-TRPA1) or HEK293 control cells (HEK) as described in the Data Supplement. Light grey areas represent SEM. (*B*) Areas under the curves were calculated and plotted as bars (green bars, HEK-TRPA1; black bars HEK). (*C-E*) Lysates from HEK293 control cells (HEK) and HEK cells stably expressing TRPA1 protein (HEK-TRPA1) were incubated in Western Blots with commercially available antibodies as described in the Data Supplement (*C*) TRPA1 (Thermo-Fisher Scientific, #PA5-22833, lot RH2256214, 1:400), (*D*) TRPA1 (Merck Millipore, #ST1685, lot J8261-6G8 1:200), (*E*) TRPA1 (Santa Cruz Biotechnology, #sc-376495, lot C2621, 1:100), (*F*) TRPA1 (Alomone labs, #ACC-037, lot ACC037AN2025, 1:1000). Beta-actin served as loading control. Data were analyzed by a Kruskal-Wallis test (*B*) and are presented as mean ± SEM (n = 3 independent experiments in B). For B, $^*P < 0.05$ versus the value in HEK control cells.

**Figure E3:** $Ca^{2+}$ imaging in primary human lung fibroblasts (HLFs) treated with the TRPA1 channel inhibitor A-967079. (*A*) HLFs were cultured with solvent (Solv.) or TGF-β1 (TGF-β1) and analyzed in $Ca^{2+}$ imaging experiments as described in Materials and Methods. Cells were stimulated with JT010 (75 nM) at the indicated time point. Light colored areas represent SEM. (*B*) Areas under the curves were calculated and plotted as bars (JT010, Solv., black bar; JT010, TGF-β1, red bar). (*C*) HLFs were cultured with solvent (Solv.) or TGF-β1 (TGF-β1) and analyzed in $Ca^{2+}$ imaging experiments as described in Materials and Methods. Cells were treated with A-967079 (500 nM) or solvent for 2 min. and stimulated with AITC (3 µM) at the indicated time points. Light colored areas represent SEM. (*D*) Areas under the curves were calculated and plotted as bars (Solv., grey bar; + TGF-β1, red bar). AITC and A-967079 (+) or solvent (-) were added as indicated. Data were analyzed by a Mann-Whitney test (*B*) or one-way ANOVA (*D*) and are presented as mean ± SEM (n = 5 independent experiments each in A-D). For B, *P < 0.05, **P < 0.01,***P < 0.001, ****P <0.0001 versus the value in cells treated with solvent alone.

**Figure E4:** Quantification of transcription and expression of fibrotic marker proteins in human lung fibroblasts (HLFs) cultured with or without TGF-β1. (*A, B*) Summaries of quantifications of α-smooth muscle actin (αSMA) (*A*) and collagen1A1 (COL1A) (*B*) protein levels in HLFs cultured with TGF-β1 and transfected with TRPA1 specific siRNAs (siRNA TRPA1) or scrambled siRNAs (siRNA Ctrl.). (*C*) α-smooth muscle actin (α-SMA) protein expression was quantified by Western blotting using a specific antiserum in HLFs cultured with or without TGF-β1 and/or treated with the TRPA1 activator JT010. Alpha-tubulin served as loading control. (*D*) Summary of α-SMA protein expression normalized to α-tubulin in HLFs cultured with or without TGF-β1 and/or treated with the TRPA1 activator JT010. (*E*) Collagen1A1 (COL1A1) protein expression was quantified by Western blotting using a specific antiserum in HLFs cultured with or without TGF-β1 and/or treated with the TRPA1 activator JT010. Alpha-tubulin served as loading control. (*F*) Summary of COL1A1 protein expression normalized to α-tubulin in HLFs cultured with or without TGF-β1 and/or treated with the TRPA1 activator JT010. (*G*) Indirect quantification of expression of plasminogen-activator-inhibitor 1 (PAI-1) activity in human lung fibroblasts (HLFs) cultured with (+)

or without (-) TGF-β1 or treated with (+) or without (-) the TRPA1-specific activator AITC. Data were analyzed by a Mann-Whitney test (*A, B*) or two-way ANOVA (*D, F, G*) and are presented as mean ± SEM (n = 3 -4 independent cell isolations each in A, B and C). For A, B, D, F, G *P < 0.05, ****P <0.0001 versus the value in cells treated with solvent alone.

**Figure E5:** Identification of MAPK p38 protein involved in TRPA1-mediated inhibition of TGF-β1-induced transcription. (*A*) Lysates from fibroblasts transfected with TRPA1 specific (siRNA TRPA1) or scrambled siRNA (siRNA Ctrl.) and treated with (+) or without (-) AITC were incubated in a Western Blot of with specific antibodies directed against phosphorylated MAPK p38 (p-MAPK p38) and MAPK p38 (MAPK p38). Alpha-tubulin served as loading control. (*B*) Summary of the p-MAPK p38/MAPK p38 quantification normalized to α-tubulin by Western Blotting in HLFs transfected with TRPA1 specific siRNAs (siTRPA1, blue bars) or scrambled siRNAs (siCtrl., grey bars) incubated with (+) or without (-) AITC. (*C*) Lysates from fibroblasts pretreated with TGF-β1, transfected with TRPA1 specific (siRNA TRPA1) or scrambled siRNA (siRNA Ctrl.) and incubated with (+) or without (-) AITC were incubated in a Western Blot of with specific antibodies directed against phosphorylated MAPK p38 (p-MAPK p38) and MAPK p38 (MAPK p38). Alpha-tubulin served as loading control. (*D*) Summary of the p-MAPK p38/MAPK p38 quantification normalized to α-tubulin by Western Blotting in HLFs pretreated with TGF-β1, transfected with TRPA1 specific siRNAs (siTRPA1, blue bars) or scrambled siRNAs (siCtrl., grey bars) and incubated with (+) or without (-) AITC. (*E*) Lysates from fibroblasts pretreated with TGF-β1, transfected with TRPA1 specific (siRNA TRPA1) or scrambled siRNA (siRNA Ctrl.) and incubated with (+) or without (-) AITC were incubated in a Western Blot of with specific antibodies directed against phosphorylated ERK1/2 (p-ERK1/2) and ERK1/2 (ERK1/2). Alpha-tubulin served as loading control. (*F*) Summary of the p-ERK1/2/ERK1/2 quantification normalized to α-tubulin and control by Western Blotting in HLFs pretreated with TGF-β1, transfected with TRPA1 specific siRNAs (siTRPA1, blue bars) or scrambled siRNAs (siCtrl., grey bars) and incubated with (+) or without (-) AITC. Data were analyzed by two-way ANOVA and are presented as mean ± SEM. Cells are from at least three independent donors (n = 4). **P < 0.01 versus the value in cells treated with solvent alone.

**Figure E6:** Identification of proteins involved in TRPA1-mediated inhibition of TGF-β1-induced transcription. (*A*) Summary of the p-SMAD2/SMAD2 quantification normalized to β-actin and control by Western Blotting in HLFs pretreated with TGF-β1, with (+) or without (-) A-967079 and incubated with (+) or without (-) AITC. (*B*) Detection of SMAD3 protein in the cytoplasmic fraction of HLFs pretreated with (+) or without (-) TGF-β1 and incubated with AITC for 0, 10 or 30 minutes. (*C*) Summary of SMAD3 protein quantifications in HLFs pretreated with (+) or without (-) TGF-β1 and incubated with AITC for 10 minutes. Data were analyzed by a Kruskal-Wallis test (*A*) or two-way ANOVA (*C*) and are presented as mean ± SEM.

**Figure E7:** Quantification of apoptosis and cell viability in human lung fibroblasts (HLFs). (*A*) HLFs cultured with solvent or TGF-β1 and transfected with a TRPA1-specific siRNA (siTRPA1) or a scrambled siRNA (siCtrl.) were incubated with the solvent (DMSO), AITC (AITC 3 µM or 6 µM) or tamoxifen (TAM) as positive control. Caspase 3 activity was detected as described in the Data Supplement. (*B*) HLFs cultured with solvent or TGF-β1 and transfected with a TRPA1-specific siRNA (siTRPA1) or a scrambled siRNA (siCtrl.) were incubated with the solvent (DMSO) or AITC for 2 or 24 h. WST assays were performed as described in the Data Supplement. Data were analyzed by two-way ANOVA and are presented as mean ± SEM (n = 3 independent cell isolations each in A-B). For A, ****P <0.0001 versus the value in cells treated with AITC or DMSO alone.

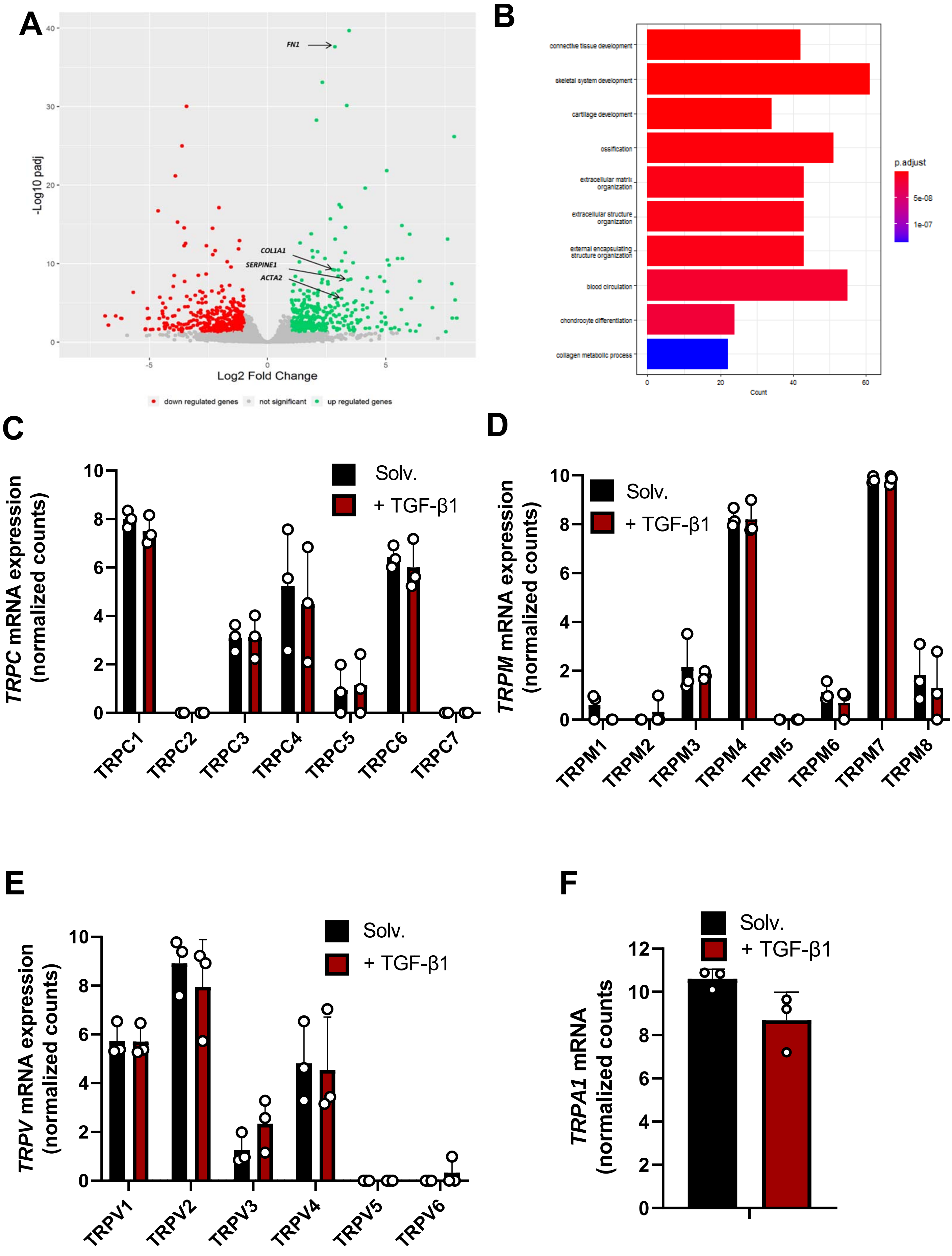
A
FN1
COL1A1
SERPINE1
ACTA2
-Log10 padj
Log2 Fold Change
down regulated genes
not significant
up regulated genes
B
connective tissue development
skeletal system development
cartilage development
ossification
extracellular matrix organization
extracellular structure organization
external encapsulating structure organization
blood circulation
chondrocyte differentiation
collagen metabolic process
Count
p.adjust
5e-08
1e-07
C
TRPC mRNA expression (normalized counts)
Solv.
+ TGF-β1
TRPC1
TRPC2
TRPC3
TRPC4
TRPC5
TRPC6
TRPC7
D
TRPM mRNA expression (normalized counts)
Solv.
+ TGF-β1
TRPM1
TRPM2
TRPM3
TRPM4
TRPM5
TRPM6
TRPM7
TRPM8
E
TRPV mRNA expression (normalized counts)
Solv.
+ TGF-β1
TRPV1
TRPV2
TRPV3
TRPV4
TRPV5
TRPV6
F
TRPA1 mRNA (normalized counts
Solv.
+ TGF-β1

**Figure E1**

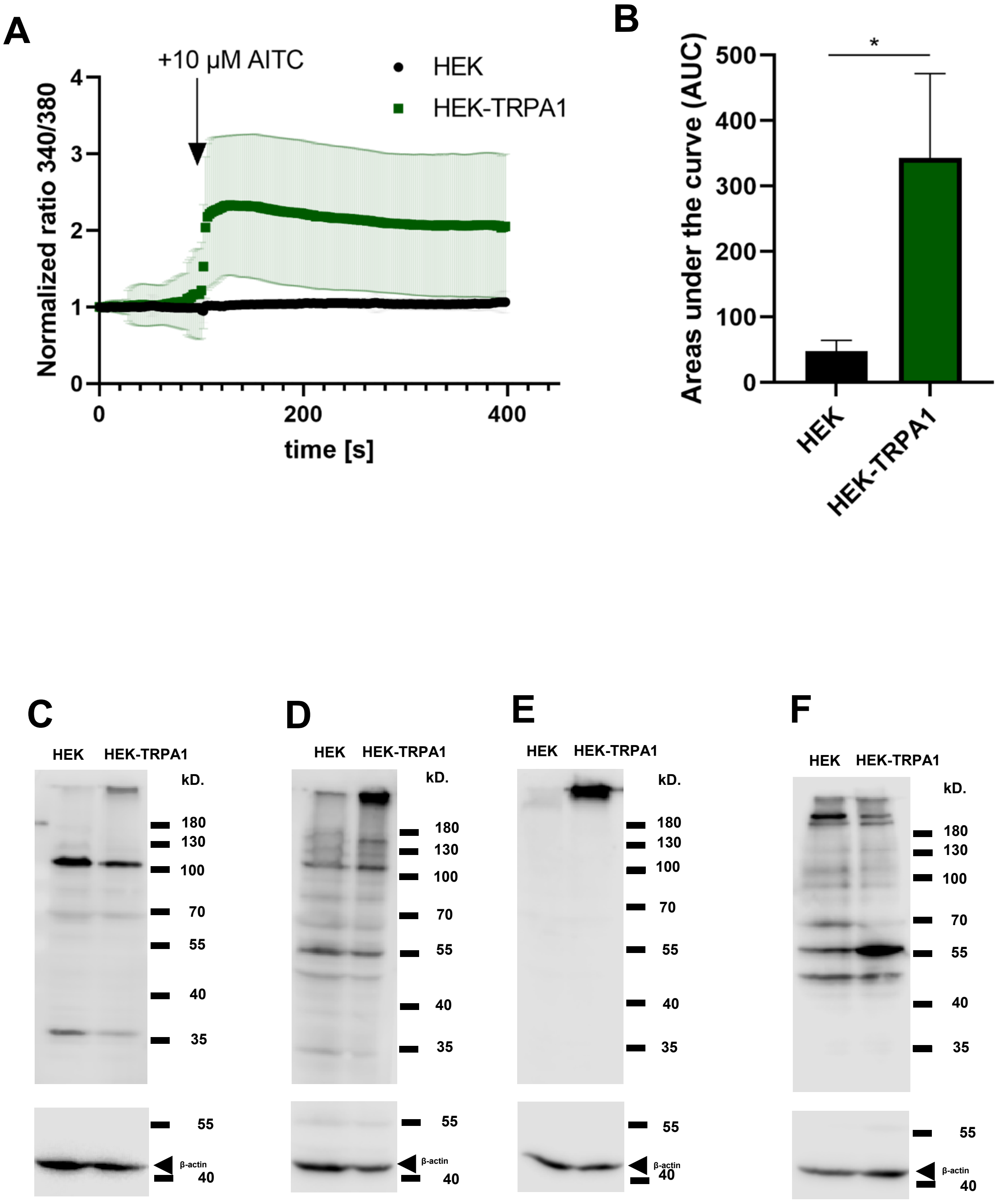
A
+10 µM AITC
HEK
HEK-TRPA1
Normalized ratio 340/380
0
1
2
3
4
0
200
400
time [s]
B
Areas under the curve (AUC)
*
0
100
200
300
400
500
HEK
HEK-TRPA1
C
HEK
HEK-TRPA1
kD.
180
130
100
70
55
40
35
55
β-actin
40
D
HEK
HEK-TRPA1
kD.
180
130
100
70
55
40
35
55
β-actin
40
E
HEK
HEK-TRPA1
kD.
180
130
100
70
55
40
35
55
β-actin
40
F
HEK
HEK-TRPA1
kD.
180
130
100
70
55
40
35
55
β-actin
40

**Figure E2**

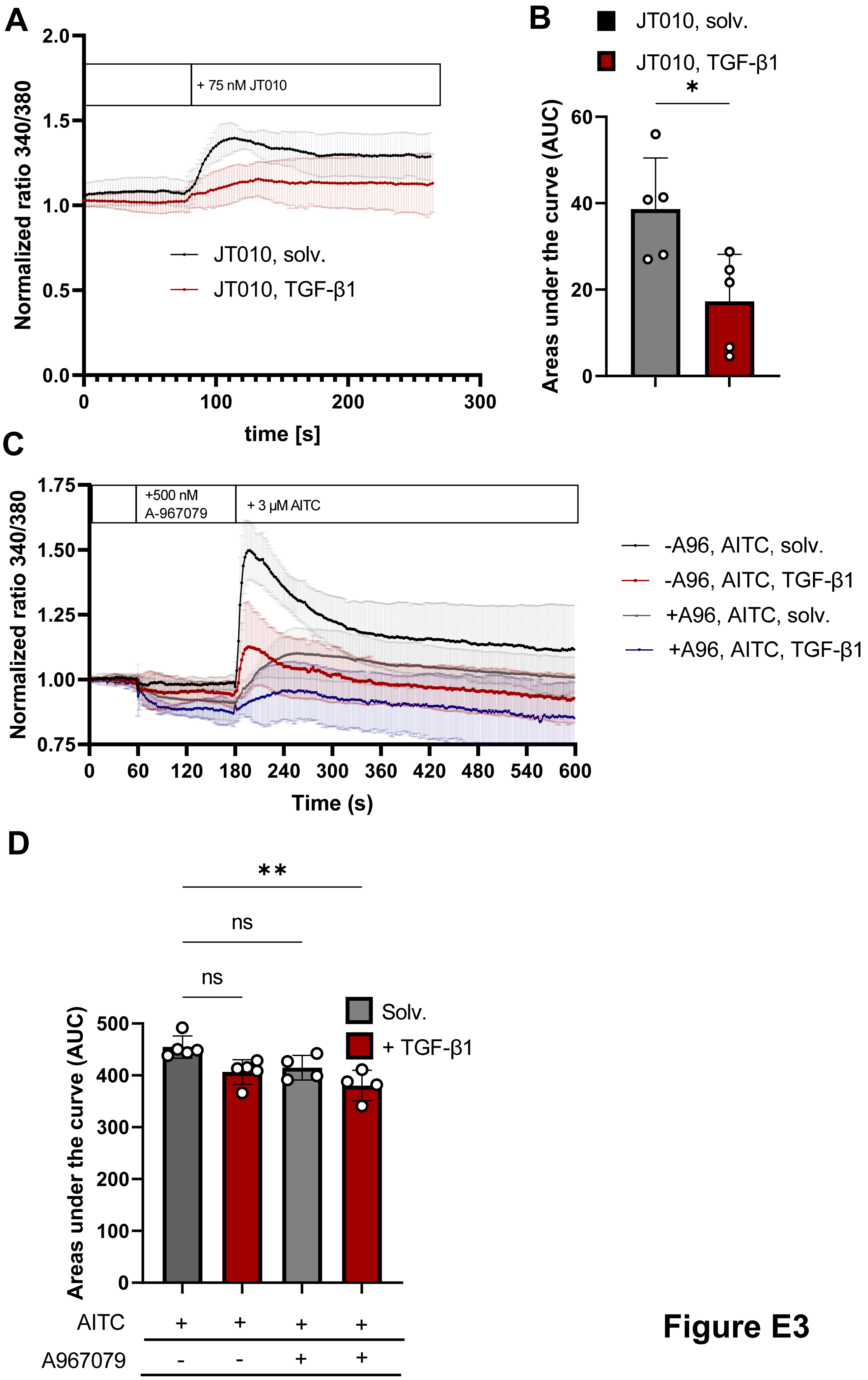
A
Normalized ratio 340/380
2.0
1.5
1.0
0.5
0.0
+ 75 nM JT010
JT010, solv.
JT010, TGF-β1
0
100
200
300
time [s]
B
JT010, solv.
JT010, TGF-β1
*
Areas under the curve (AUC)
60
40
20
0
C
Normalized ratio 340/380
1.75
1.50
1.25
1.00
0.75
+500 nM A-967079
+ 3 µM AITC
-A96, AITC, solv.
-A96, AITC, TGF-β1
+A96, AITC, solv.
+A96, AITC, TGF-β1
0
60
120
180
240
300
360
420
480
540
600
Time (s)
D
**
ns
ns
Areas under the curve (AUC)
500
400
300
200
100
0
Solv.
+ TGF-β1
AITC + + + +
A967079 - - + +

**Figure E3**

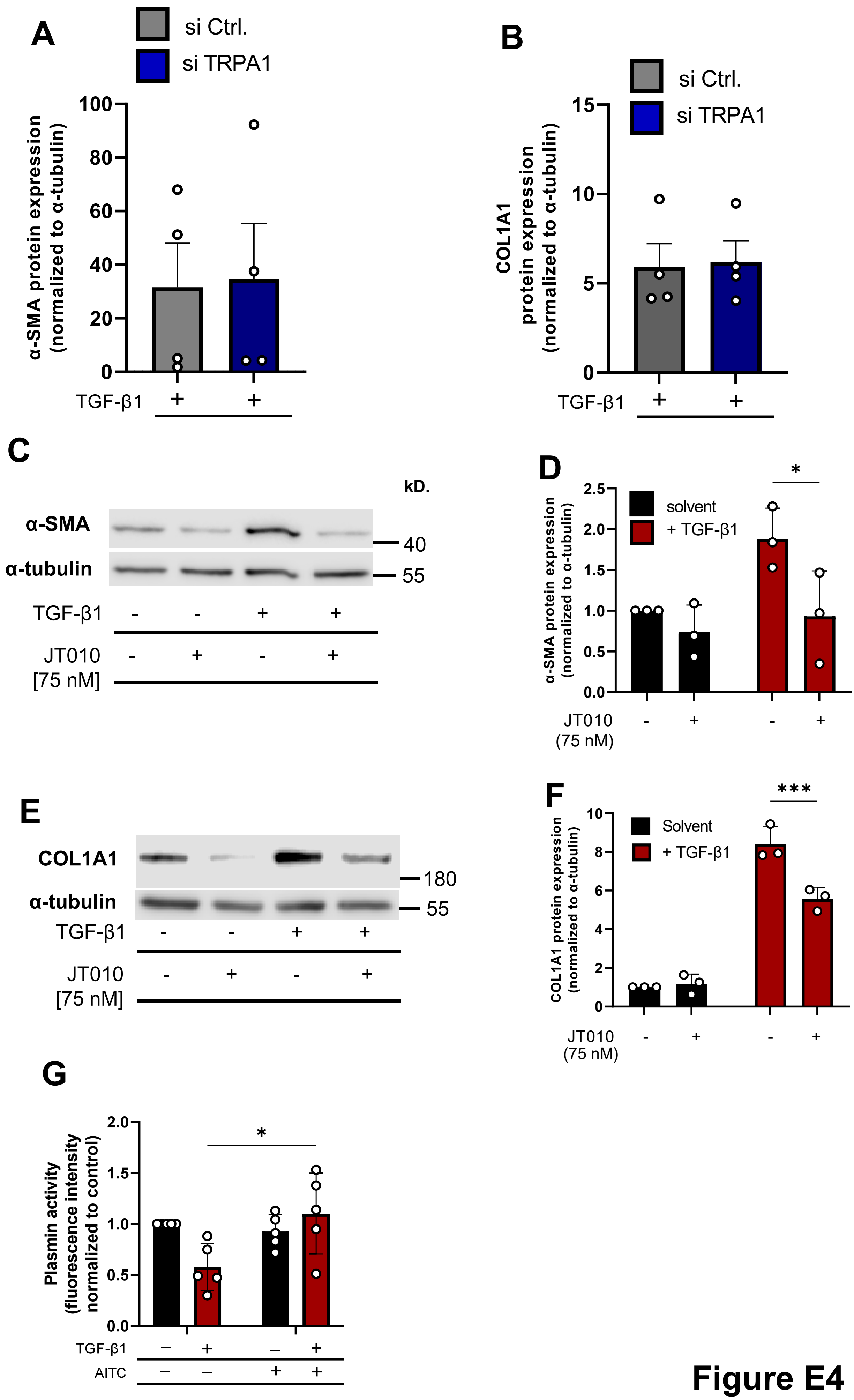
A
si Ctrl.
si TRPA1
α-SMA protein expression (normalized to α-tubulin)
TGF-β1 + +
B
si Ctrl.
si TRPA1
COL1A1 protein expression (normalized to α-tubulin)
TGF-β1 + +
C
kD.
α-SMA
α-tubulin
40
55
TGF-β1 - - + +
JT010 [75 nM] - + - +
D
*
solvent
+ TGF-β1
α-SMA protein expression (normalized to α-tubulin)
JT010 (75 nM) - + - +
E
COL1A1
α-tubulin
180
55
TGF-β1 - - + +
JT010 [75 nM] - + - +
F
***
Solvent
+ TGF-β1
COL1A1 protein expression (normalized to α-tubulin)
JT010 (75 nM) - + - +
G
*
Plasmin activity (fluorescence intensity normalized to control)
TGF-β1 – + – +
AITC – – + +

Figure E4

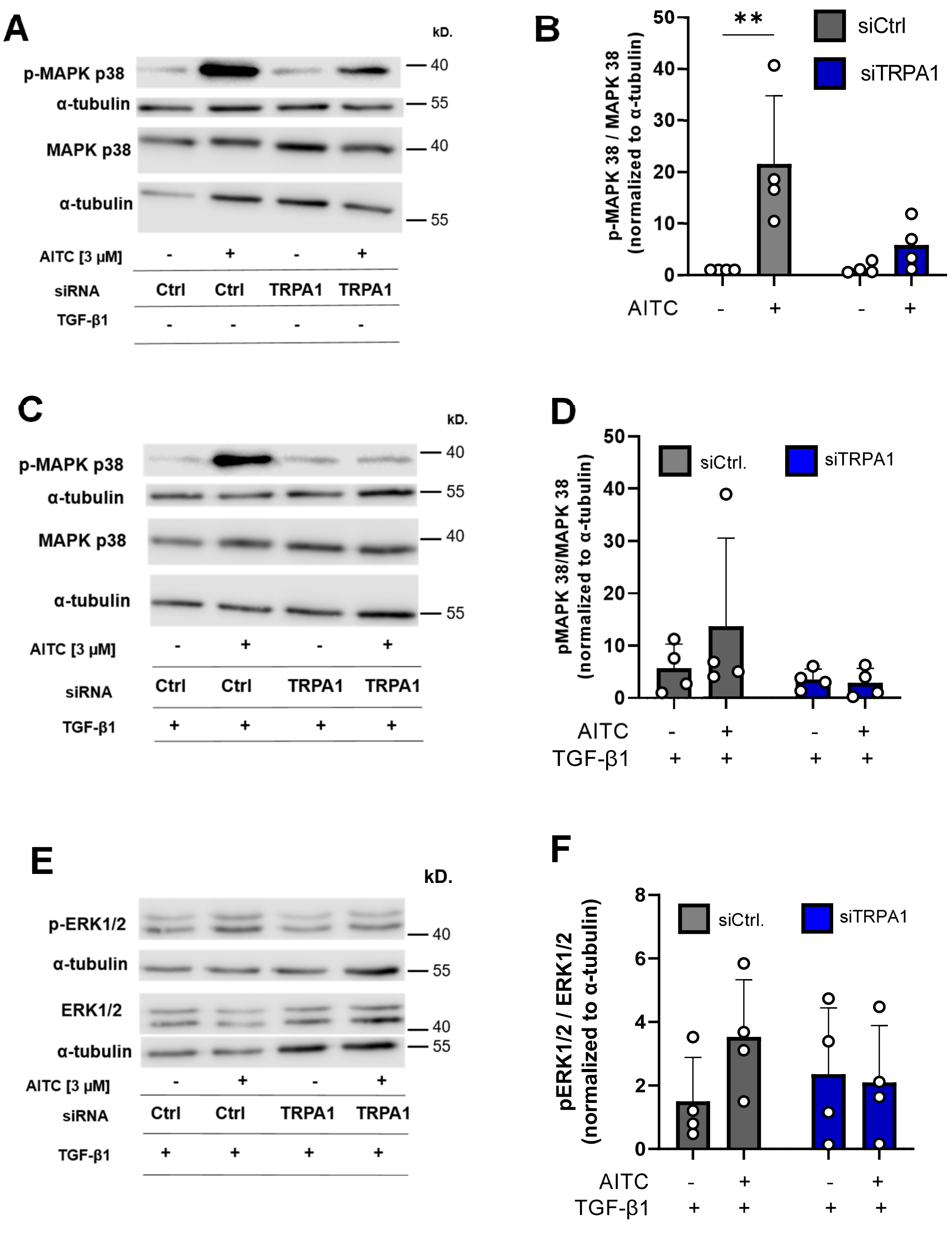
A
kD.
p-MAPK p38
α-tubulin
MAPK p38
α-tubulin
40
55
40
55
AITC [3 µM] - + - +
siRNA Ctrl Ctrl TRPA1 TRPA1
TGF-β1 - - - -
B
p-MAPK 38 / MAPK 38 (normalized to α-tubulin)
**
siCtrl
siTRPA1
AITC - + - +
C
kD.
p-MAPK p38
α-tubulin
MAPK p38
α-tubulin
AITC [3 µM] - + - +
siRNA Ctrl Ctrl TRPA1 TRPA1
TGF-β1 + + + +
D
pMAPK 38/MAPK 38 (normalized to α-tubulin)
siCtrl.
siTRPA1
AITC - + - +
TGF-β1 + + + +
E
kD.
p-ERK1/2
α-tubulin
ERK1/2
α-tubulin
AITC [3 µM] - + - +
siRNA Ctrl Ctrl TRPA1 TRPA1
TGF-β1 + + + +
F
pERK1/2 / ERK1/2 (normalized to α-tubulin)
siCtrl.
siTRPA1
AITC - + - +
TGF-β1 + + + +

**Figure E5**

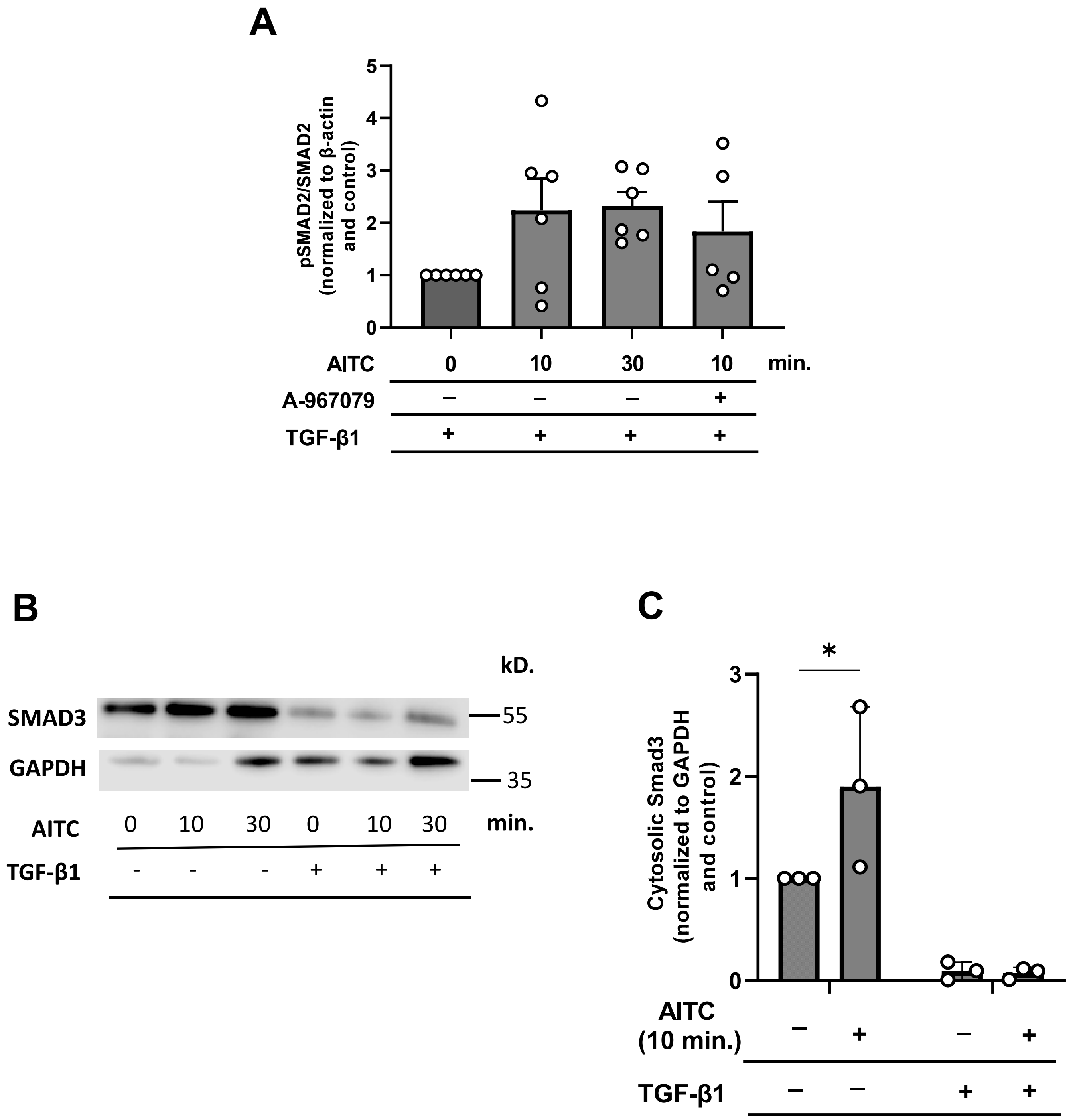
A
pSMAD2/SMAD2 (normalized to β-actin and control)
0
1
2
3
4
5
AITC
0
10
30
10
min.
A-967079
–
–
–
+
TGF-β1
+
+
+
+
B
kD.
SMAD3
55
GAPDH
35
AITC
0
10
30
0
10
30
min.
TGF-β1
-
-
-
+
+
+
C
*
Cytosolic Smad3 (normalized to GAPDH and control)
0
1
2
3
AITC (10 min.)
–
+
–
+
TGF-β1
–
–
+
+

Figure E6

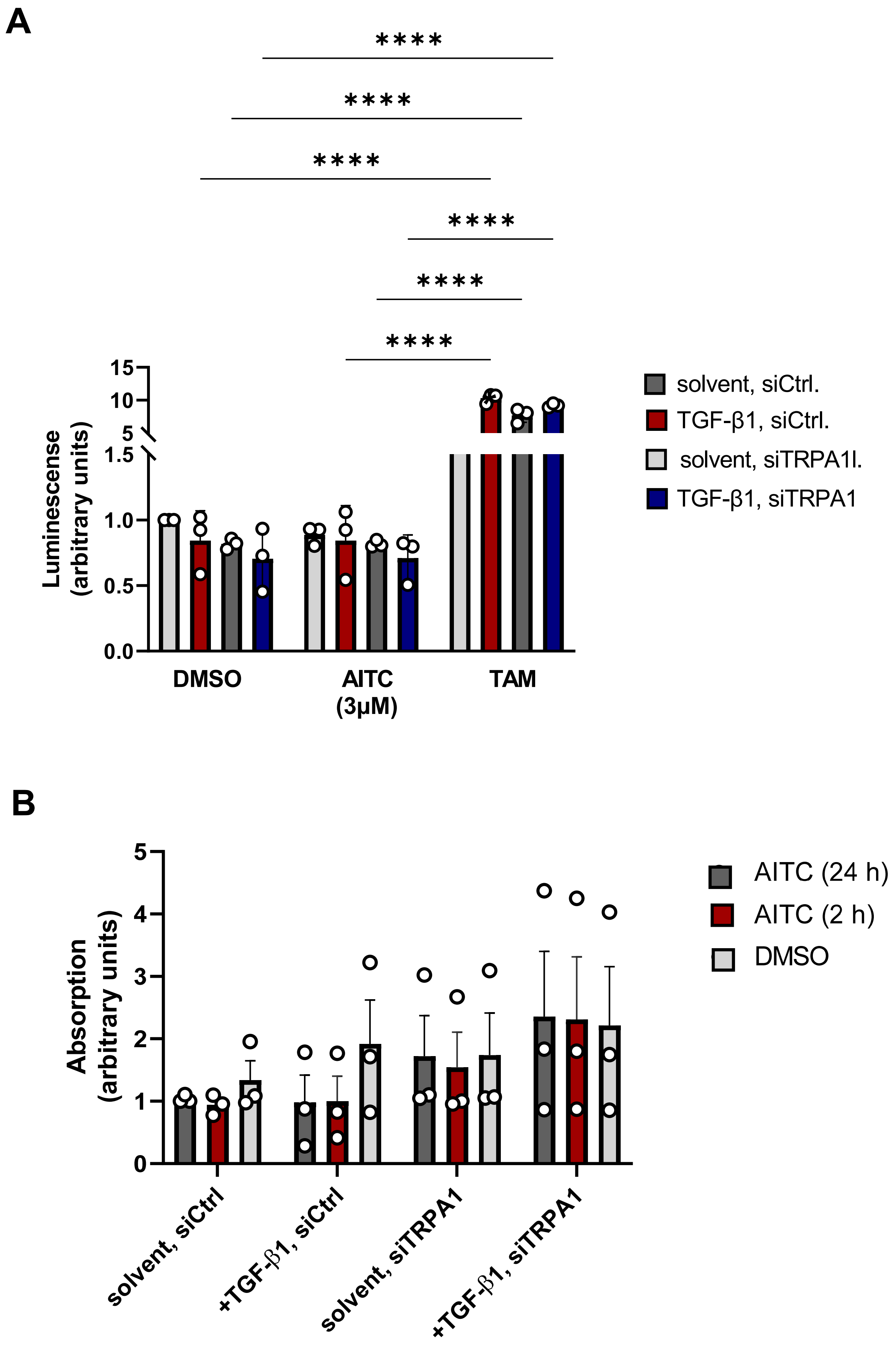
A
****
****
****
****
****
****
15
10
5
1.5
1.0
0.5
0.0
Luminescense
(arbitrary units)
DMSO
AITC
(3µM)
TAM
solvent, siCtrl.
TGF-β1, siCtrl.
solvent, siTRPA1l.
TGF-β1, siTRPA1
B
5
4
3
2
1
0
Absorption
(arbitrary units)
solvent, siCtrl
+TGF-β1, siCtrl
solvent, siTRPA1
+TGF-β1, siTRPA1
AITC (24 h)
AITC (2 h)
DMSO

**Figure E7**
